# A Role for Synaptonemal Complex in Meiotic Mismatch Repair

**DOI:** 10.1101/2021.05.28.446251

**Authors:** Karen Voelkel-Meiman, Ashwini Oke, Arden Feil, Alexander Shames, Jennifer Fung, Amy MacQueen

## Abstract

During meiosis a large subset of interhomolog recombination repair intermediates form within the physical context of the synaptonemal complex (SC), a protein-rich structure assembled at the interface of aligned homologous chromosomes. However, the functional relationship between SC structure and homologous recombination remains poorly defined. In prior work we determined that tripartite SC is dispensable for recombination in *S. cerevisiae*; SC central element proteins Ecm11 and Gmc2 instead limit the number of recombination events. Here we report that while dispensable for recombination *per se*, SC central element proteins influence the processing of interhomolog recombination intermediates in a manner that minimizes errors in mismatch correction. Failure to correct mis-paired bases within heteroduplex at meiotic recombination sites leads to genotypically sectored colonies (post meiotic segregation events) arising from mitotic proliferation of mismatch-containing spores. We discovered an increase in post-meiotic segregation at the *THR1* locus in cells lacking Ecm11 or Gmc2, or in the SC-deficient but crossover recombination-proficient *zip1[Δ21-163]* mutant. High-throughput sequencing of octad meiotic products revealed a genome-wide increase in recombination events with uncorrected mismatches in *ecm11* mutants relative to wild type. Meiotic cells missing Ecm11 also display longer gene conversion tracts, but tract length alone does not account for the higher frequency of uncorrected mismatches. Interestingly, the per-nucleotide mismatch frequency is elevated in *ecm11* mutants when analyzing all gene conversion tracts, but is similar between wild type and *ecm11* if one considers only those events with uncorrected mismatches. Our data suggest that a subset of recombination events is similarly susceptible to mismatch repair errors in both wild type and *ecm11* strains, but in *ecm11* mutants many more recombination events fall into this inefficient repair category. Finally, we observe elevated post-meiotic segregation at *THR1* in mutants with a dual deficiency in MutSγ-mediated crossover recombination and SC assembly, but not in the *mlh3* mutant, which lacks MutSγ crossovers but has abundant SC. We propose that SC structure promotes efficient mismatch repair of joint molecule recombination intermediates resolved via both MutSγ-associated and MutSγ-independent pathways, and is the molecular basis for elevated post-meiotic segregation in both MutSγ crossover-proficient (*ecm11, gmc2*) and MutSγ crossover-deficient (*msh4, zip3*) strains.

## Introduction

While discouraged in vegetative cells, reciprocal recombination between homologous chromosomes is a critical feature of meiosis. In conjunction with sister chromatid cohesion, such crossover recombination events provide transient attachments between chromosomes that enable partner homologs to orient and disjoin toward opposite poles of the meiosis I spindle. Crossover-fated recombination intermediates physically associate with a widely conserved structure called the synaptonemal complex (SC), a ∼100 nm wide protein-rich assembly found at the interface of lengthwise-aligned homologous chromosome axes during meiotic prophase (Page and Hawley 2004). Particularly in budding yeast where SC is dispensable for crossover formation, (Voelkel-Meiman *et al*. 2015; Voelkel-Meiman *et al*. 2016; Voelkel-Meiman *et al*. 2019) the functional relationship between assembled SC and interhomolog recombination remains obscure.

Meiotic recombination initiates with Spo11-mediated DNA double strand breaks (DSBs), and is completed via one of several overlapping pathways that involve single-stranded DNA degradation, strand invasion of intact homologous DNA duplex, and new DNA synthesis (reviewed in Hunter 2015). Joint molecule recombination intermediates arising from these processing events are then guided through one of at least two mechanistically-distinct pathways to produce a noncrossover (resulting in DNA duplexes that have not undergone reciprocal exchange) or a crossover. In Synthesis Dependent Strand Annealing (SDSA), unligated joint molecule intermediates are dismantled without endonuclease activity. In Double Strand Break Repair (DSBR), post-strand invasion DNA synthesis ultimately leads to a ligated joint molecule carrying two Holliday junctions. A prominent class of Holliday junction intermediates in *S. cerevisiae* - those that rely on the MutSγ complex for their stable formation - are typically processed via a MutLγ endonuclease pathway that mediates biased, asymmetrical cleavage of the joint molecule (non-crossed DNA strands are cleaved at one Holliday junction while crossed strands are cleaved at the other junction), resulting in reciprocal (crossover) exchange (Argueso *et al*. 2004; Zakharyevich *et al*. 2010; de Muyt *et al*. 2012; Zakharyevich *et al*. 2012; Kulkarni *et al*. 2020). On the other hand, MutLγ-independent resolution pathways - such as the one involving the structure-selective endonucleases Mus81-Mms4 together with Yen1 - are associated with unbiased cleavage of the two Holliday junctions, which results in the formation of both noncrossovers and crossovers (de Los Santos *et al*. 2001; Argueso *et al*. 2004; de Muyt *et al*. 2012).

The SC is poised to play a functional role in the formation or processing of Holliday junction recombination intermediates in particular. SC is fully assembled along the entire interface of aligned chromosome axes during mid-meiotic prophase (Zickler and Kleckner 1998; Zickler and Kleckner 1999), when ligated joint molecule intermediates are most abundant. By contrast, the majority of noncrossover events in budding yeast are thought to arise via an SDSA mechanism relatively early in meiotic prophase (Allers and Lichten 2001a), prior to full SC assembly. Furthermore, the formation of a substantial fraction of joint molecules relies on pro-crossover proteins that are also required for timely SC assembly (Lynn *et al*. 2007) and localize as discrete foci within the central region of the SC (Voelkel-Meiman *et al*. 2019). Such factors include the MutSγ complex (consisting of a heterodimer of Msh4 and Msh5, eukaryotic homologs of the bacterial MutS mismatch repair protein) (Novak *et al*. 2000; Novak *et al*. 2001), which is thought to directly stabilize Holliday junction DNA structures (Snowden *et al*. 2004), the Mer3 helicase (Nakagawa and Ogawa 1999), the XPF-XRCC-related Zip4-Zip2-Spo16 subcomplex (de Muyt *et al*. 2018), the E3 SUMO ligase Zip3 (Agarwal and Roeder 2000; Cheng *et al*. 2006; Tsubouchi *et al*. 2008), as well as the SC structural protein, Zip1 (Storlazzi *et al*. 1996; Voelkel-Meiman *et al*. 2016; Voelkel-Meiman *et al*. 2019). Finally, the resolution of MutSγ Holliday junction intermediates in *S. cerevisiae* is dependent on the Polo-like Cdc5 kinase, which also triggers disassembly of SC (Allers and Lichten 2001a; Sourirajan and Lichten 2008).

From an ultrastructural view, the ∼100 nanometer wide SC structure consists of three major parts. Within the context of SC, parallel-aligned, protein-rich axes of fully replicated meiotic chromosomes are referred to as lateral elements. Lateral elements are bridged along their lengths by rod-like, transverse filament proteins which are oriented perpendicular to the long axis of the aligned chromosomes, like the rungs of a ladder. Each “rung” of the SC ladder is comprised of two transverse filament protein complexes oriented with their N termini toward the midline of the SC structure. Finally, central element proteins assemble at the midline of the SC structure near the N termini of transverse filament proteins (Page and Hawley 2004). Zip1 is the only known transverse filament protein of the budding yeast SC (Sym *et al*. 1993; Dong and Roeder 2000), while Ecm11 and Gmc2 contribute to the assembly of SC central element (Humphryes *et al*. 2013; Voelkel-Meiman *et al*. 2013). The central region of the budding yeast SC is dynamic, exhibiting continuous deposition of transverse filament and central element proteins during prophase even after primary full-length structures have assembled (Voelkel-Meiman *et al*. 2012; Voelkel-Meiman *et al*. 2013). This means that during meiotic prophase SC structures have the capacity to acquire progressively more central region in between homologous axes.

Despite its physical proximity to recombination intermediates, however, tripartite SC is dispensable for meiotic recombination in budding yeast. Meiotic cells that lack the SC central element proteins Ecm11 or Gmc2, or that express a version of the Zip1 protein missing most of its N terminus, fail to assemble SC but exhibit an increased abundance of MutSγ crossovers (Voelkel-Meiman *et al*. 2016). The budding yeast SC structure thus plays a role in attenuating interhomolog recombination events, possibly by discouraging the accumulation of additional DNA DSBs, and/or by preventing nascent recombination intermediates from engaging with the homolog (versus the sister chromatid) for repair (Subramanian *et al*. 2016; Mu *et al*. 2020).

While the SC is dispensable for crossover recombination in budding yeast, genetic data from our prior study suggests that SC central element proteins nevertheless do influence the processing of MutSγ intermediates. Meiotic cells missing the MutLγ component, Mlh3, ordinarily exhibit the same degree of diminished crossover recombination as mutants lacking the MutSγ component, Msh4. However, this is not the case when the SC central element protein Ecm11 is missing: *ecm11 mlh3* double mutants show a substantially larger number of crossovers than *ecm11 msh4* double mutants (Voelkel-Meiman *et al*. 2016). Thus, the presence of budding yeast SC structure (or SC central element proteins) dictates the degree to which MutSγ recombination intermediates can resolve to a crossover outcome.

Here we report additional evidence that budding yeast SC influences the processing of interhomolog recombination intermediates, namely the capacity of such intermediates to undergo successful mismatch repair. DNA mismatch repair is central to meiotic recombination because the heteroduplex that forms as a result of strand invasion and DNA synthesis involves homologous but not necessarily identical DNA strands (Holliday 1964; White *et al*. 1985; Allers and Lichten 2001b). The formation of interhomolog recombination intermediates during meiosis results in heteroduplex containing mismatched bases, owing to sequence differences between homologous partner chromosomes. Such mismatches are normally corrected by mismatch repair (MMR) enzymes, in a direction leading either to restoration of original sequence or conversion to the sequence of the partner DNA duplex (reviewed in Borts *et al*. 2000). Specific eukaryotic homologs of bacterial mismatch repair enzymes have been shown to have a major role in mismatch repair during meiosis, including the MutS family member Msh2 and the MutL family members Pms1 and Mlh1 (Williamson *et al*. 1985; Kramer *et al*. 1989; Prolla *et al*. 1994b). A molecular picture for how and when recombination intermediates interface with mismatch repair enzymes during meiosis is unclear. Interestingly, however, data reported in (Getz *et al*. 2008) led the authors to propose that mismatch repair is a stronger feature of the MutSγ-mediated recombination pathway relative to MutSγ-independent recombination in budding yeast meiotic cells.

Here we report that mutants missing structural components of the SC exhibit diminished meiotic mismatch repair. In one such mutant missing the Ecm11 protein, we show that an increased number of recombination sites genome-wide display uncorrected DNA mismatches. We furthermore observe inefficient mismatch repair in SC-deficient meiotic cells that have either lost or gained MutSγ crossovers, arguing against a simple correlation between MutSγ-mediated recombination and robust mismatch repair. Our findings lead us to propose that the SC structure directly affects the processing of DNA joint molecule intermediates in a manner that ensures successful repair of mismatched nucleotides.

## Results

### Diminished meiotic mismatch repair at *THR1* in the absence of SC central element proteins

When mismatch repair fails during meiosis in budding yeast, spore products inherit mismatch-carrying DNA duplexes (Figure 1A). Daughter cells resulting from the first mitotic division of such spores inherit two distinct genotypes, and additional vegetative growth results in the formation of a sectored colony consisting of cells of either genotype. If the two genotypes are linked to an observable phenotype, these post-meiotic segregation (PMS) events result in phenotypically sectored colonies.

**Figure 1.**
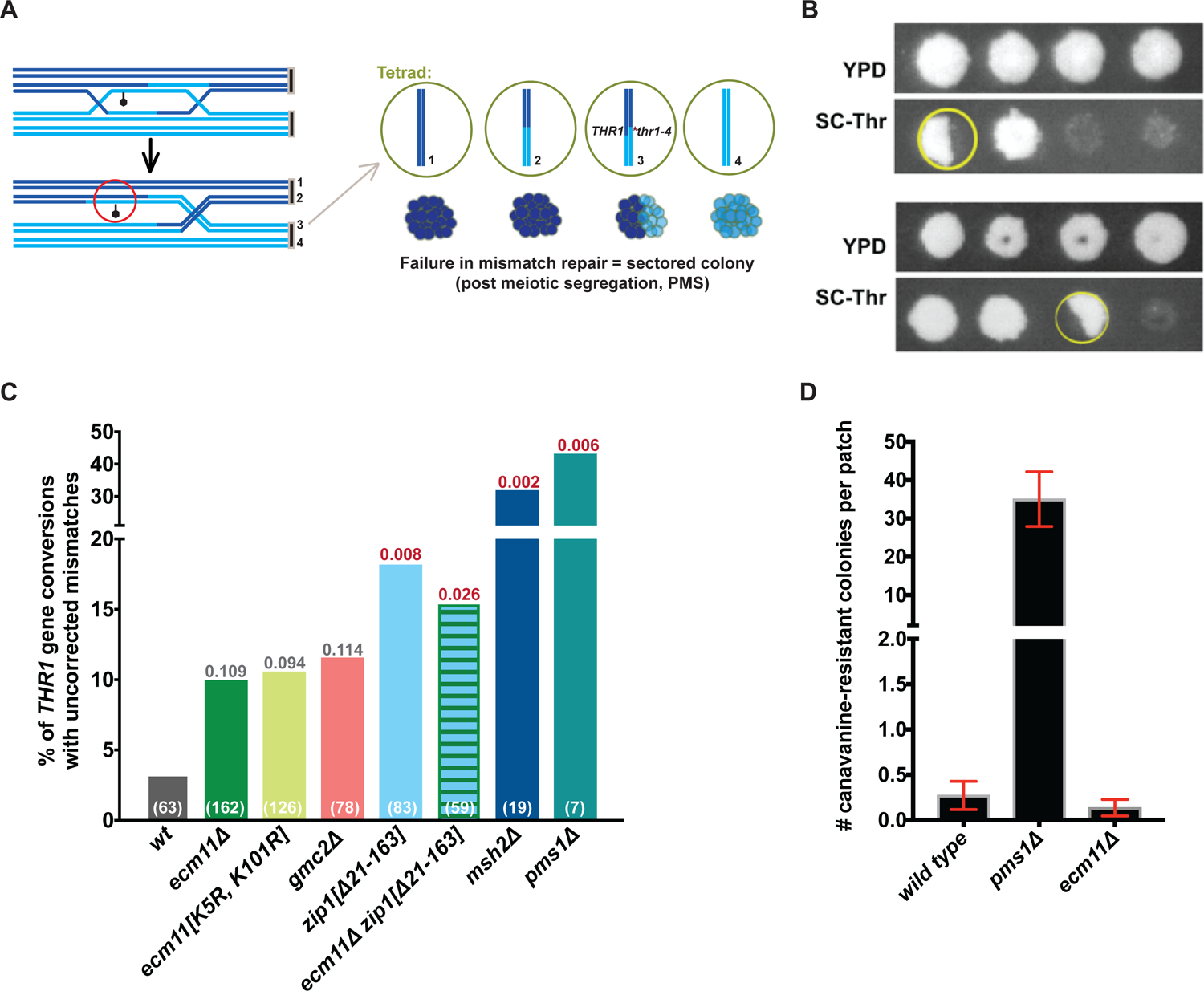
Increased meiotic mismatch repair errors in the absence of synaptonemal complex central element. **A**) Cartoon illustrates the role of mismatch repair during meiotic recombination. A failure to correct mismatches (black lollipop) in heteroduplex DNA can result in post-meiotic segregation (PMS) events, corresponding to a genotypically (and potentially phenotypically) sectored colony within a tetrad. **B**) Two examples of 4-spore viable tetrads grown on rich media (upper panels) or SC-Thr dropout media (lower panels). Tetrads derive from diploids heterozygous for the *thr1-4* point mutation, which corresponds to an AàT transversion at nucleotide 22 of the *THR1* open reading frame. The yellow circled colony is phenotypically sectored for growth on SC-Thr media. **C**) Graph plots the percentage of *THR1* gene conversion-associated tetrads (tetrads exhibiting non-mendelian segregation at *THR1*) that exhibit a phenotypically sectored colony on SC-Thr, reflecting the occurrence during meiosis of an uncorrected DNA mismatch. The total number of tetrads (meioses) observed with non-mendelian segregation of the *THR1* alleles is indicated (white) at the bottom of the corresponding bar for each strain. For example, 63 gene conversion events at *THR1* were identified among 1163 four-spore viable tetrads from our wild-type strain, and two (3.2%) of these tetrads displayed a colony with sectored growth on SC-Thr. *P* values (derived from a Fisher’s Exact Test comparing mutant data to wild type) are indicated above individual columns; values in red are considered statistically significant, while those in grey not quite significant. See Table 1 for additional features of post-meiotic segregation events for 4-spore viable and non-4-spore viable tetrads from these strains. **D**) Bar graph indicates the average median of canavanine-resistant colonies per patch of wild-type, *pms1* or *ecm11* cells measured in four independent forward-mutation-to-canavanine-resistance experiments involving eleven patches each (see Methods). The range of medians is indicated by error bars: The median number of papillae per patch in each *pms1* experiment was 8, 38, 42, and 43; the median number of papillae per patch in each of the four wild-type and four *ecm11* experiments was zero.

**Table 1.** Errors in meiotic recombination-associated DNA mismatch repair at THR1 in SC-deficient mutants. *(Relates to Figure 1)* The tables in (**A**) and (**B)** list meiotic segregation data for *THR1* in strains heterozygous for several genetic marker alleles, including *thr1-4*. The first three data columns in (**A**) give the percentage of four-spore viable tetrads exhibiting gene conversion (non 2:2 segregation events) at *THR1*, which is an approximate measure of total interhomolog recombination frequency at this locus. The next two data columns indicate the number and percentage, among total interhomolog recombination events at *THR1*, of four-spore viable tetrads displaying a sectored colony on SC-Thr media. A sectored colony (post-meiotic segregation event, or PMS) indicates the failure to repair a mismatch within heteroduplex associated with a *THR1* gene conversion tract. The subsequent columns list the number of tetrads observed for each strain that fall into each possible category of non-mendelian segregation; category names reflect the proportion of DNA single strands (8 DNA strands represent a given set of four chromatids comprising two homologs) encoding one versus the other *THR1* allele. 6:2 or 2:6, and 8:0 or 0:8 reflect one or two conventional gene conversion events, respectively, at *THR1* within a tetrad, with no sectored colonies apparent. By contrast, 5:3 or 3:5, and 7:1 or 1:7 tetrads display one sectored colony, while Ab6:2 and Ab4:4 tetrads each display two sectored colonies. **B**) The number of one-, two-, or three-spore viable meiotic products with a colony that is sectored for growth on SC-Thr (column 1). Subsequent columns indicate features of the post-meiotic segregation event (note ratios indicate whether the event occurred in a one-spore viable (1:1), a two-spore viable (3:1, Ab2:2, 1:3) or a three-spore viable (5:1, 3:3, Ab4:2, 1:5) meiosis. See Table S4 for full strain genotypes.

| A |  | Nonmendelian Segregation @ *THR1* in 4-spore Viable Tetrads |  |  |  |  |  |  |  |  |  |  |  |  |  |  |
| --- | --- | --- | --- | --- | --- | --- | --- | --- | --- | --- | --- | --- | --- | --- | --- | --- |
|  |  | # 4-SV<br>TETRADES | # NON 2:2<br>TETRADES | % NON<br>2:2 | # TETRADES<br>W/PMS | % NON 2:2<br>W/PMS | *no mismatch* |  |  |  | *uncorrected mismatch (PMS)* |  |  |  |  |  |
| GENOTYPE (STRAIN) |  |  |  |  |  |  | 8:0 | 6:2 | 2:6 | 0:8 | 7:1 | 5:3 | Ab<br>6:2 | Ab<br>4:4 | 3:5 | 1:7 |
| *WT*<br>(K842) |  | 1163 | 63 | 5.4 | 2 | 3.2 | 0 | 40 | 21 | 0 | 0 | 0 | 0 | 0 | 1 | 1 |
| *ecm11Δ*<br>(K857) |  | 944 | 162 | 17.2 | 16 | 9.9 | 1 | 76 | 67 | 2 | 2 | 3 | 1 | 0 | 9 | 1 |
| *ecm11[K5R,K101R]*<br>(K1123/K1124) |  | 528 | 126 | 23.9 | 13 | 10.3 | 2 | 64 | 45 | 2 | 0 | 6 | 0 | 0 | 6 | 1 |
| *gmc2Δ*<br>(K906) |  | 511 | 78 | 15.3 | 9 | 11.5 | 1 | 35 | 32 | 1 | 1 | 3 | 0 | 1 | 4 | 0 |
| *zip1[Δ21-163]*<br>(AF6) |  | 612 | 83 | 13.6 | 15 | 18.1 | 0 | 36 | 32 | 0 | 0 | 6 | 0 | 0 | 9 | 0 |
| *zip1[Δ21-163] ecm11Δ*<br>(AF5) |  | 521 | 59 | 11.3 | 9 | 15.3 | 0 | 32 | 18 | 0 | 0 | 4 | 0 | 0 | 5 | 0 |
| *msh2Δ*<br>(AM3697/AM4003) |  | 219 | 19 | 8.7 | 6 | 31.6 | 0 | 6 | 7 | 0 | 1 | 3 | 0 | 0 | 1 | 1 |
| *pms1Δ*<br>(AM3696/AM4004) |  | 117 | 7 | 6.0 | 3 | 42.9 | 0 | 4 | 0 | 0 | 0 | 3 | 0 | 0 | 0 | 0 |

**Table 1. Errors in meiotic recombination-associated DNA mismatch repair at *THR1* in SC-deficient mutants.**
| B | Uncorrected Mismatches (PMS) @ *THR1* in NON-4-Spore Viable Meiotic Products |  |  |  |  |  |  |  |  |  |
| --- | --- | --- | --- | --- | --- | --- | --- | --- | --- | --- |
|  | GENOTYPE (STRAIN) | # PMS<br>NON-4 SV | 5:1 | 3:1 | 3:3 | 1:1 | Ab<br>4:2 | Ab<br>2:2 | 1:3 | 1:5 |
|  | *WT*<br>(K842) | 0 | 0 | 0 | 0 | 0 | 0 | 0 | 0 | 0 |
|  | *ecm11Δ*<br>(K857) | 4 | 1 | 2 | 0 | 0 | 0 | 0 | 0 | 1 |
|  | *ecm11[K5R,K101R]*<br>(K1123/K1124) | 3 | 0 | 1 | 1 | 0 | 0 | 0 | 0 | 1 |
|  | *gmc2Δ*<br>(K906) | 3 | 0 | 0 | 3 | 0 | 0 | 0 | 0 | 0 |
|  | *zip1[Δ21-163]*<br>(AF6) | 3 | 0 | 0 | 3 | 0 | 0 | 0 | 0 | 0 |
|  | *zip1[Δ21-163] ecm11Δ*<br>(AF5) | 9 | 0 | 0 | 5 | 0 | 1 | 1 | 0 | 0 |
|  | *msh2Δ*<br>(AM3697/AM4003) | 6 | 1 | 1 | 3 | 0 | 0 | 0 | 0 | 1 |
|  | *pms1Δ*<br>(AM3696/AM4004) | 0 | 0 | 0 | 0 | 0 | 0 | 0 | 0 | 0 |

We were surprised to discover an elevated frequency of PMS events at the *THR1* locus in SC central element-deficient mutants (*ecm11*, *gmc2*, *zip1[Δ21-163]*) heterozygous for *thr1-4* (Figure 1B, C, Table 1). The *thr1-4* allele has a single point mutation at nucleotide position 22 within the *THR1* open reading frame. In evaluating interhomolog recombination frequency in SC-deficient meiotic mutants, we expected to observe conventional gene conversion events at *THR1*; such tetrads are classified primarily as 6:2 and 2:6 events, where three out of four DNA duplexes comprising the two homologous sets of sister chromatids carry the genomic sequence associated either *thr1-4* or *THR1*. However, we also encountered non-conventional gene conversion events only rarely observed in the wild type strain. In four spore viable tetrads classified as carrying 5:3, 3:5, 7:1, or 1:7 events, the Watson and Crick strands of one DNA duplex encode different *THR1* alleles; the spore with this mismatch-carrying chromosome produces a sectored colony on SC-Thr media. In 1163 four-spore viable tetrads from wild-type meiotic cells, 63 gene conversion events were observed at the *THR1* locus, and only two of these were non-conventional (3:5 and 1:7). The frequency of post-meiotic segregation (uncorrected DNA mismatches) involving *THR1* in our wild-type strain is thus 3.2% (Figure 1C, Table 1A). In 944 four-spore viable tetrads from meiotic cells missing the SC central element component Ecm11, 162 interhomolog gene conversion events were observed at the *THR1* locus (the higher frequency of interhomolog recombination at *THR1* is consistent with the established role SC central element proteins in limiting interhomolog recombination (Voelkel-Meiman *et al*. 2016)), and sixteen (9.9%) of these correspond to non-conventional events (Figure 1C, Table 1A). Similarly, 15.1% of 86 *THR1* gene conversions in the *ecm11[K5R, K101R]* mutant (which produces a nonSUMOylatable version of Ecm11 that does not support SC assembly (Humphryes *et al*. 2013)), and nine of the 78 observed *THR1* gene conversions (11.5%) in *gmc2* null mutants, were observed to be non-conventional (Figure 1C, Table 1A). Using a Fishers Exact Test, the approximately three to five fold increase in post-meiotic segregation at *THR1* displayed by SC central element-deficient mutants is not strongly statistically significant. However, the consistent phenotype among the three SC-deficient mutants, including the presence of additional sectored colony events in non-four spore viable meiotic products in the mutants (Table 1B), suggests a role for SC central element proteins in ensuring accurate mismatch repair during meiosis.

*ZIP1* encodes an SC transverse filament protein whose N terminus colocalizes with SC central element proteins. Strains carrying the *zip1[Δ21-163]* allele produce a Zip1 protein missing most of its predicted unstructured N terminus and, similar to *ecm11* and *gmc2* mutants, fail to assemble SC although they retain high levels of MutSγ recombination. We found that *zip1[Δ21-163]* strains exhibit a statistically significant increase (relative to wild type) in *THR1* post-meiotic segregation: 18.1% (15/83) of detected *THR1* gene conversions are non-conventional (Figure 1C, Table 1). Similarly, 15.3% (9/59) of *THR1* gene conversion events in *ecm11 zip1[Δ21-163]* double mutants exhibit post-meiotic segregation. The mismatch repair defect displayed by SC central element-deficient mutants is less dramatic than the one associated with mutants missing the core mismatch repair proteins Msh2 or Pms1 (which respectively exhibited 32% and 43% post-meiotic segregation frequencies at *THR1* in our assay; Figure 1C, Table 1).

In mitotically dividing cell cultures, we observed an elevated frequency of mutation at the *CAN1* gene when Pms1 protein is missing but not when Ecm11 is deleted (Figure 1D). This indicates, as expected, that the mismatch repair phenotype of SC-deficient mutants is specific to meiosis. Taken together, the similar meiotic mismatch repair phenotype of five SC-deficient, MutSγ recombination-proficient strains points to a role for SC central element proteins in ensuring successful mismatch correction of heteroduplex recombination intermediates, at least at the *THR1* locus.

### Octad Rec-Seq reveals that the SC central element curtails both majority and minority interhomolog recombination events

To determine whether errors in mismatch repair displayed by SC central element deficient mutants are unique to the *THR1* locus, we carried out high-throughput sequence analysis of meiotic products from wild type, *ecm11*, *msh2* and *pms1* YJM789/S96 hybrid diploid strains (see Methods), which are heterozygous for ∼60,000 single nucleotide polymorphisms (SnPs) (Anderson *et al*. 2011; Mancera *et al*. 2011; Oke *et al*. 2014). We isolated genomic DNA from individual members of an octad derived from the four meiotic spore products of each hybrid strain; octads were isolated by separating mother and daughter descendants immediately following the first mitotic division of each spore within a four-spore-viable tetrad (Figure 2A, see Methods). Genomic DNA for each mother or daughter cell within the octad was then processed for Illumina sequencing, and the resulting sequences were analyzed to trace parental origins of DNA across the entire genome of each meiotic product, revealing recombination signatures.

**Figure 2.**
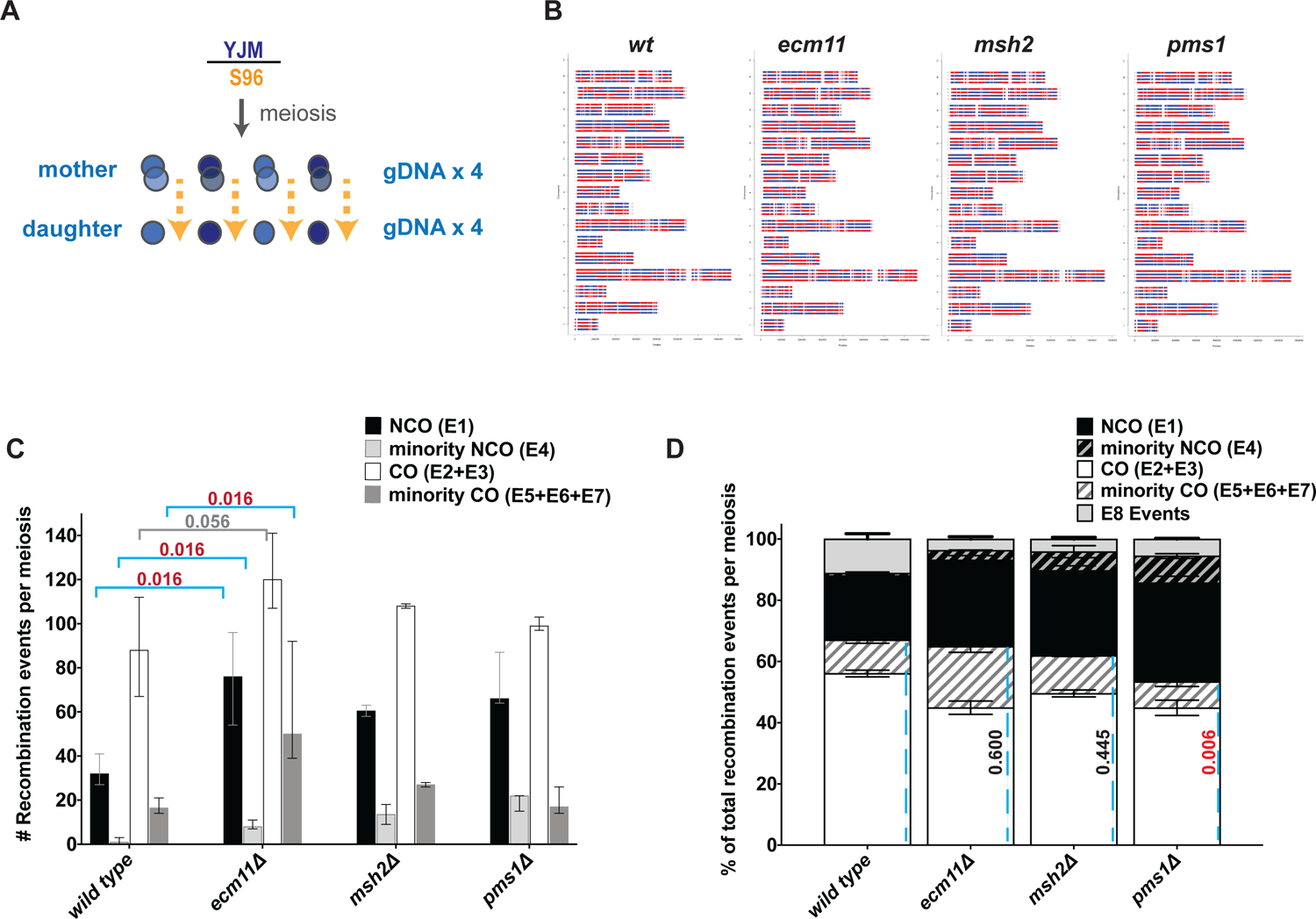
Genome-wide sequence analysis of *ecm11* meiotic products indicates a role for SC central element in limiting both majority and minority interhomolog recombination events. **A**) Cartoon illustrates the Octad Rec-Seq experiment performed on meiotic products of hybrid diploids generated by crossing YJM789 and YJMS96 genetic backgrounds (Anderson *et al*. 2011; Mancera *et al*. 2011; Oke *et al*. 2014). ∼60,000 single nucleotide polymorphisms exist across the genome in the YJM789/S96 hybrid (Anderson *et al*. 2011; Oke *et al*. 2014). Tetrads were isolated from hybrids induced to undergo meiosis, and after the first mitotic division mother and daughter cells were physically separated. Genomic DNA was isolated from each of the four mother and four daughter members of the octad and prepared for Illumina sequencing. (**B**) Genome-wide recombination maps generated by Octad Rec-Seq for each YJM789/S96 hybrid strain (wild type, *ecm11*, *msh2*, or *pms1* homozygote). Colors indicate YJM789 or S96 DNA sequence for each of the four chromatids of every yeast chromosome (stacked along the *y* axis), giving a rough impression of crossover exchange frequency in a given meiosis. One tetrad example for each genotype is shown. The bar graph in (**C**) plots the median number and range of interhomolog recombination events detected by Octad-RecSeq in four wild-type, five *ecm11*, two *msh2* and three *pms1* meioses (strains indicated on the *x* axis). Recombination values were obtained from either the “mother” or the “daughter” tetrad component of a given octad; in cases where the numbers did not precisely match between mother and daughter tetrad, the largest value was selected. Recombination events are classified as “E1”, “E2”, “E3”, “E4”, “E5”, “E5A”, “E6”, “E7”, or “E8” based on DNA sequence signatures, as described in (Oke *et al*. 2014); see Figure S1 for diagrammatic cartoons, and last section of this legend for narrative description of classes. A Mann Whitney Test determined that the averaged total noncrossover values (E1+E4) and averaged total crossover (E2+E3+E5+E6+E7) values obtained from the five *ecm11* meiotic datasets are significantly increased relative to wild type (*P* = 0.016 for each group), but these values in the *msh2* and *pms1* datasets are not significantly different from wild type (*P* = 0.133, 0.057 for noncrossovers and *P* = 0.133, 0.400 for crossovers respectively). Similarly, the majority noncrossover (E1), minority noncrossover (E4), and the combined minority crossover (E5+E6+E7) averaged values for five *ecm11* meioses are significantly greater than the corresponding averaged values of the wild type datasets (two-tailed *P* values indicated in red on the graph). In contrast, the average number of majority crossover (E2+E3) events in *ecm11*, while larger, is not significantly different from wild type (*P* = 0.056; indicated in grey on the graph). The majority and minority noncrossover and (crossover values in *msh2* and *pms1* are not significantly different from wild type (*P* =0.133 and 0.057 for *msh2* and *pms1* E1 values, *P* = 0.067 and 0.057 for *msh2* and *pms1* E4 values, *P* = 0.533 and 0.400 for *msh2* and *pms1* E2+E3 values, and *P* = 0.133 and 0.971 for *msh2* and *pms1* E5+E6+E7 values). The bar graph in (**D**) shows the proportion of total recombination events per meiosis represented by each noncrossover and crossover class. The mean and standard error of the mean values for four wild type, five *ecm11*, two *msh2* and three *pms1* octads are plotted. A Fisher’s Exact Test found no significant difference in the average proportion of total crossovers (blue dotted line) among interhomolog events in *ecm11* and *msh2* meioses compared to wild type (*P* = 0.600, 0.445, respectively), although the proportions of majority and minority classes of crossover events do vary significantly; see Table 2 and Figure S1. Interestingly, the average proportion of crossovers among total interhomolog recombination events in *pms1* meioses was found to be significantly decreased compared to wild type (*P* = 0.006). Note chromosome 7 recombination events for two wild-type octads were excluded from these analyses, due to chromosome 7 disomy. <u>Recombination signatures</u>: Meiotic crossovers and noncrossovers form through pathways that involve strand invasion followed by DNA synthesis. NCOs predominantly arise via an “SDSA” mechanism in budding yeast, where strand invasion and new DNA synthesis is followed by dismantling of the joint molecule before its ligation. A small fraction of wild type noncrossovers may also arise from unbiased resolution of ligated joint molecules. Recombination events with “E1” or “E4” DNA signatures are classified as noncrossover events. The E1 event makes up the majority noncrossover class based on its abundance in wild type strains; this event displays a 3:1 tract involving one chromatid and is not within 5 kb of another interhomolog event. An E4 event corresponds to a discontinuous noncrossover event, where two or more 3:1 tracts arrange in tandem on one chromatid with regions of 2:2 marker segregation between. Most crossovers arise from biased cutting of the two junctions in a double Holliday junction. E2 and E3 events represent the majority of crossovers in wild type strains, and are the simplest expected result from biased cutting of the double Holliday junction. E2 events correspond to the presence of two crossed-over chromatids with or without detectable gene conversion (GC) tracts and not within 5 kb of another interhomolog event. E3 events also represent simple crossovers but are associated with a discontinuous gene conversion tract on one of the recombined chromatids. E5, E5A, E6, and E7 events correspond to minority DNA sequence signatures generated by joint molecule processing events that are uncommon in wild type. E5 events correspond to two or more apparent exchange events within 5 kb of each other and involving only two chromatids (an apparent double crossover event, although E5 events could reflect noncrossovers formed via symmetrical resolution of a joint molecule). E6 recombination signatures reflect two or more interhomolog events within 5 kb of each other and involving three chromatids. E7 signatures correspond to two or more interhomolog events within 5 kb of each other and involving four chromatids. E8 events correspond to 4:0 conversion events, which may reflect a pre-meiotic gene conversion. Note that E8 events are not included in either consolidated noncrossover or crossover class percentages (instead are only included when individual subcategory data is presented).

**Table 2.** Genome-wide frequencies of meiotic interhomolog recombination events identified by Octad Rec-Seq. *(Relates to Figures 2, S1)* The table lists the raw number and percentages of interhomolog recombination events detected by Octad-RecSeq in four wild-type, five *ecm11*, two *msh2* and three *pms1* meioses (strains and individual octad identifier indicated in the left column). Recombination values were obtained from either the “mother” or “daughter” tetrad component of a given octad; in cases where the numbers did not precisely match between mother and daughter tetrad, the largest value was selected. Recombination events are classified as “E1”, “E2”, “E3”, “E4”, “E5”, “E5A”, “E6”, “E7”, or “E8” based on DNA sequence signatures, as described in Figure 2 legend and (Oke *et al*. 2014). Data are plotted in Figures 2 and S1 graphs. Note chromosome 7 recombination events for two wild-type octads were excluded from these analyses, due to chromosome 7 disomy.

|  |  | Noncrossover |  |  |  | Crossover |  |  |  |  |  |  |  |  |  |  |  |  |  |  |
| --- | --- | --- | --- | --- | --- | --- | --- | --- | --- | --- | --- | --- | --- | --- | --- | --- | --- | --- | --- | --- |
|  | total IH<br>events<br>(E1-E8) | #E1 | %E1 | #E4 | %E4 | #E2 | %E2 | #E3 | %E3 | #E5+E5A | %E5 | #E6 | %E6 | #E7 | %E7 | % NCO<br>(E1+E4) | %<br>majority<br>CO<br>(E2+E3) | %<br>minority<br>CO<br>E5+E6+E7 | #E8 | %E8 |
| wt octad 1 no ch7 | 148 | 34 | 23.0 | 0 | 0.0 | 73 | 49.3 | 10 | 6.8 | 10 | 6.8 | 5 | 3.4 | 3 | 2.0 | 23.0 | 56.1 | 12.2 | 13 | 8.8 |
| wt octad 4 no ch7 | 125 | 27 | 21.6 | 1 | 0.8 | 61 | 48.8 | 6 | 4.8 | 4 | 3.2 | 10 | 8.0 | 0 | 0.0 | 22.4 | 53.6 | 11.2 | 16 | 12.8 |
| wt octad 7 | 167 | 30 | 18.0 | 3 | 1.8 | 76 | 45.5 | 17 | 10.2 | 12 | 7.2 | 7 | 4.2 | 2 | 1.2 | 19.8 | 55.7 | 12.6 | 20 | 12.0 |
| wt octad 8 | 190 | 41 | 21.6 | 1 | 0.5 | 96 | 50.5 | 16 | 8.4 | 3 | 1.6 | 10 | 5.3 | 2 | 1.1 | 22.1 | 58.9 | 7.9 | 21 | 11.1 |
| em11 octad 9 | 249 | 69 | 27.7 | 8 | 3.2 | 85 | 34.1 | 39 | 15.7 | 9 | 3.6 | 25 | 10.0 | 6 | 2.4 | 30.9 | 49.8 | 16.1 | 8 | 3.2 |
| ecm11 octad 10 | 355 | 96 | 27.0 | 11 | 3.1 | 99 | 27.9 | 42 | 11.8 | 34 | 9.6 | 48 | 13.5 | 10 | 2.8 | 30.1 | 39.7 | 25.9 | 15 | 4.2 |
| ecm11 octad 11 | 226 | 54 | 23.9 | 8 | 3.5 | 85 | 37.6 | 22 | 9.7 | 17 | 7.5 | 29 | 12.8 | 4 | 1.8 | 27.4 | 47.3 | 22.1 | 7 | 3.1 |
| ecm11 octad 12 | 270 | 87 | 32.2 | 8 | 3.0 | 74 | 27.4 | 33 | 12.2 | 22 | 8.1 | 30 | 11.1 | 3 | 1.1 | 35.2 | 39.6 | 20.4 | 13 | 4.8 |
| ecm11 octad 13 | 250 | 76 | 30.4 | 7 | 2.8 | 89 | 35.6 | 31 | 12.4 | 15 | 6.0 | 23 | 9.2 | 1 | 0.4 | 33.2 | 48.0 | 15.6 | 8 | 3.2 |
| msh2 octad 15 | 221 | 58 | 26.2 | 18 | 8.1 | 68 | 30.8 | 39 | 17.6 | 19 | 8.6 | 7 | 3.2 | 2 | 0.9 | 34.4 | 48.4 | 12.7 | 10 | 4.5 |
| msh2 octad 16 | 215 | 63 | 29.3 | 9 | 4.2 | 69 | 32.1 | 40 | 18.6 | 12 | 5.6 | 9 | 4.2 | 5 | 2.3 | 33.5 | 50.7 | 12.1 | 8 | 3.7 |
| pms1 octad 17 | 224 | 66 | 29.5 | 22 | 9.8 | 54 | 24.1 | 43 | 19.2 | 5 | 2.2 | 21 | 9.4 | 0 | 0.0 | 39.3 | 43.3 | 11.6 | 13 | 5.8 |
| pms1 octad 18 | 207 | 64 | 30.9 | 15 | 7.2 | 64 | 30.9 | 39 | 18.8 | 2 | 1.0 | 9 | 4.3 | 3 | 1.4 | 38.2 | 49.8 | 6.8 | 11 | 5.3 |
| pms1 octad 19 | 238 | 87 | 36.6 | 22 | 9.2 | 66 | 27.7 | 33 | 13.9 | 9 | 3.8 | 6 | 2.5 | 2 | 0.8 | 45.8 | 41.6 | 7.1 | 13 | 5.5 |

First, the genomic sequence of each of the four mothers or four daughters of a given octad were analyzed to determine the frequency, position, and DNA sequence features of interhomolog recombination events genome-wide (Figure 2B, Table 2). In four wild-type octad datasets 148, 125, 167, and 190 interhomolog recombination events were detected, while 249, 355, 226, 270 and 250 events were detected in five *ecm11* octad datasets (Table 2), consistent with the increase in interhomolog recombination previously-reported for *ecm11* meiotic cells (Voelkel-Meiman *et al*. 2016). *msh2* and *pms1* octads also exhibited a slight increase in interhomolog recombination relative to wild-type: two *msh2* octads displayed 221 and 215 interhomolog events while 224, 207, and 238 events were found in three *pms1* octads (Table 2). While the increase in the total number of noncrossover and crossover recombination events in *ecm11* strains relative to wild type is significant (Two-tailed *P* value using a Mann-Whitney test = 0.016 for total noncrossover and total crossover values), the total number of interhomolog recombination events is not significantly different between wild-type strains and *msh2* or *pms1* (Figure 2B, Table 2).

Recombination events can be classified by distinct DNA sequence signatures of the four chromatids encompassing a meiotic tetrad; such subclasses that are thought to reflect distinct recombination intermediates and/or resolution mechanisms (Anderson *et al*. 2011; Oke *et al*. 2014; see Figure 2 legend). Closer analysis of recombination event subcategories in our Octad-RecSeq strains revealed that the fraction of recombination events represented by a majority crossover class for wild type (E2, representing simple asymmetric cutting of a double Holliday junction with a continuous or no detectable gene conversion tract) is significantly diminished in *ecm11*, *msh2*, and *pms1* relative to wild type (Table 2, Figure S1). A compensatory increase in mostly E3 crossover events, defined as simple crossovers with discontinuous gene conversion tracts, is observed in all three mutant strains, but the increase is significant only for *msh2* and *pms1* (Table 2, Figure S1). Interestingly, the fraction of a single minority class (E6) of crossovers that involves three chromatids is significantly increased in *ecm11* relative to all other strains analyzed (Table 2, Figure S1). Given the mismatch repair phenotype reported below for *ecm11*, the elevation in crossovers with discontinuous tracts (E3) in *ecm11, msh2* and *pms1* meioses could share a common mechanistic basis involving mismatch repair. However, the increase in events involving more than one template (E6) appears to be due to the loss of an activity belonging to Ecm11 that is not shared by Msh2 or Pms1.

Notably, while total interhomolog recombination is elevated in *ecm11* mutants, the proportion of crossovers among total interhomolog events is not significantly different from wild type (Fishers Exact Test *P* = 0.600; Figure 2D), consistent with our prior genetic analysis of noncrossover:crossover ratios at individual loci in *ecm11* (Voelkel-Meiman *et al*. 2016). By contrast with *ecm11*, and unlike the results of a prior genome-wide study (Martini *et al*. 2011), in our hybrid strains *msh2* showed no significant difference from wild type in the proportion of total events that resolve to a crossover outcome (*P* = 0.445; Figure 2D). However, *pms1* hybrids displayed a significant reduction in the proportion of total events that resolve to a crossover, relative to wild type (*P* = 0.006; Figure 2D). Using an independent genetic analysis in an isogenic, BR genetic background, we found that neither *msh2* nor *pms1* mutants display a significant decrease in genetic map length across chromosome III and much of chromosome VIII (Table S1). Thus strain background differences likely underlie the variable crossover frequency observed in mutants lacking core mismatch repair machinery.

### “Octad Rec-Seq” indicates a role for an SC structural protein in promoting efficient mismatch repair genome-wide

If DNA mismatches arising from heteroduplex at recombination sites are properly corrected during meiosis, then each pair of the four mother - daughter pairs in an octad should share precisely the same genomic sequence. On the other hand, errors in meiotic mismatch repair within gene conversion tracts of interhomolog recombination events are detectable as genomic sequence differences between mother and daughter members of a pair. Our Octad Rec-Seq analysis revealed that approximately 10% of wild-type meiotic interhomolog recombination events contain an uncorrected mismatch within the associated gene conversion tract (12, 15, 20, and 16 events in four wild-type octads; Figure 3A dark bars, Table 3), consistent with prior studies (Mancera *et al*. 2011; Martini *et al*. 2011). By contrast, a significant increase in the proportion of interhomolog recombination events with an uncorrected mismatch was found in the *ecm11* hybrid: 120, 189, 99, 97, and 66 events in five *ecm11* octads (26-53% of the total events) carry an uncorrected mismatch (*P* <0.0001; Figure 3A dark bars, Table 3). Thus, the increase in post-meiotic segregation observed genetically at the *THR1* locus in *ecm11* mutants reflects a genome-wide meiotic mismatch repair defect. As expected, a majority of recombination events detected in *msh2* and *pms1* Octad Rec-Seq datasets exhibit uncorrected mismatches: 165 and 170 recombination events (75-79% of the total events) carry an uncorrected mismatch in two *msh2* octads while 102, 163 and 194 recombination events (46-82% of total events) carry an uncorrected mismatch in three *pms1* octads, respectively (Figure 3A dark bars, Table 3).

**Figure 3.**
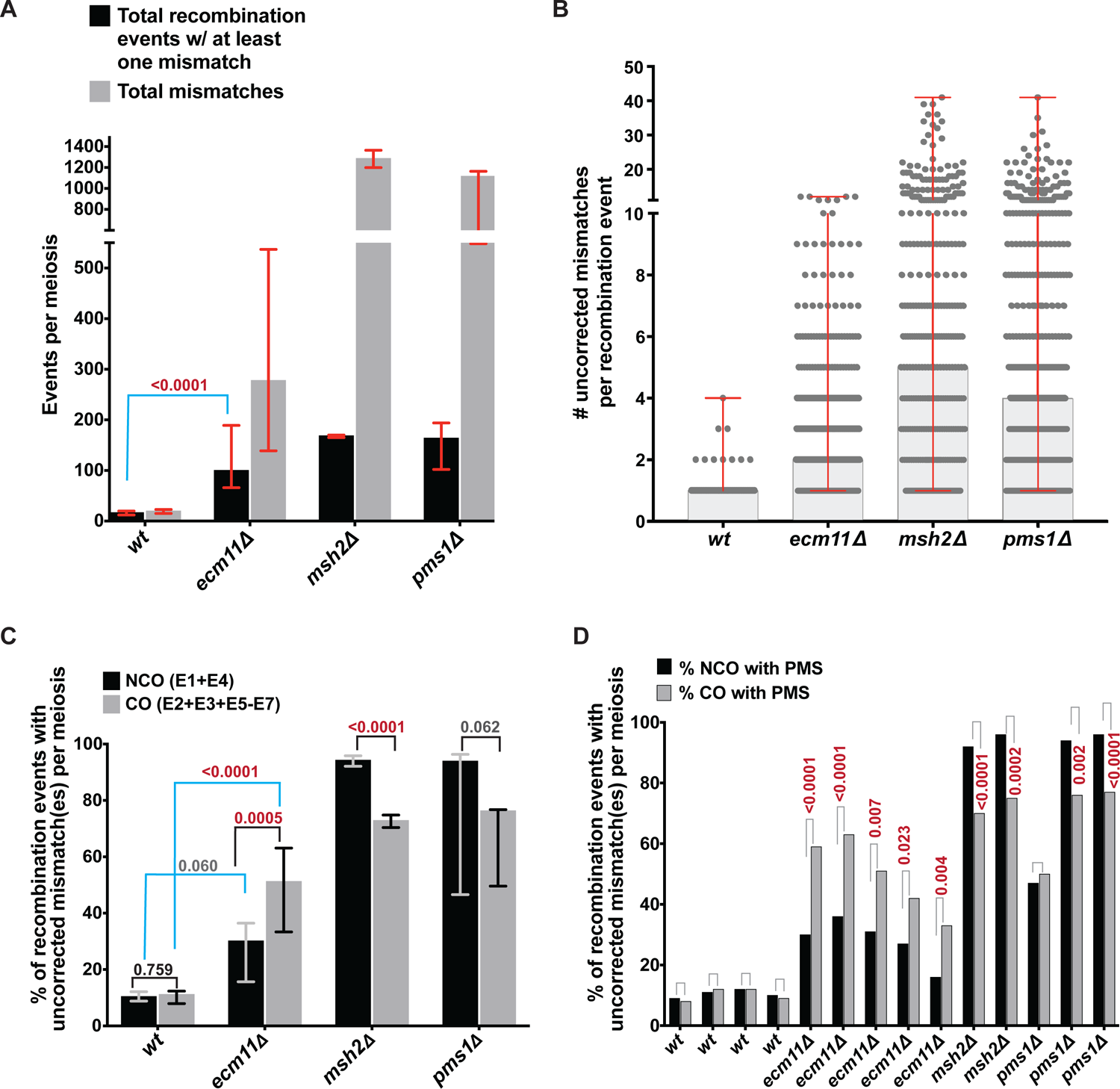
*ecm11* mutants display increased errors in meiotic mismatch repair genome-wide, particularly at crossover events. Bar graph in (**A)** plots the number of recombination events per meiosis (per octad) that carry at least one uncorrected mismatch in the associated gene conversion tract (dark bars), and the total number of uncorrected mismatches detected genome-wide (light bars). The median and range of values for four wild-type, five *ecm11*, two *msh2* and three *pms1* octads (indicated on the *x* axis) are given. The average proportion of total recombination events associated with an uncorrected mismatch is significantly increased in *ecm11*, *msh2* and *pms1* octad datasets relative to wild type datasets (*P* values for each strain determined by Fisher’s Exact Test = < 0.0001; indicated on graph for *ecm11* only). Each circle on the scatterplot in (**B**) represents a recombination event with at least one uncorrected mismatch. The *y* axis indicates the number of uncorrected mismatches detected at a given recombination site. Data from four wild-type, five *ecm11*, two *msh2* and three *pms1* meioses (Octad Rec-Seq datasets) is plotted. Red bars indicate the range, and the shaded column indicates the median number of uncorrected mismatches detected per poorly-repaired recombination site for each strain. Note that while the number of uncorrected mismatches per recombination event in *ecm11* has a larger range than wild-type, between 1 and 12, most events carry only one or two uncorrected mismatches thus the medians of *ecm11* and wild type are not dramatically different. Column graphs in (**C**) and (**D**) plot the percentage of noncrossover (dark bars) or crossover (light bars) events that carry at least one uncorrected mismatch within the associated gene conversion tract for each of the four wild-type, five *ecm11*, two *msh2* and three *pms1* Octad Rec-Seq datasets. The average median and range of values for each genotype is plotted in (**C**), while values for each individual octad are plotted in (**D**). Two-tailed *P* values are indicated above compared groups where averaged means from octad datasets were assessed for significant differences using a Fisher’s Exact Test. Blue bars indicate comparisons between *ecm11* and wild-type; a significant difference (red numbers) was found between the average proportion of crossover events with at least one uncorrected mismatch in *ecm11* mutants relative to wild-type, but the increase in average proportion of noncrossovers with an uncorrected mismatch in *ecm11* mutants relative to wild-type is not quite significant (grey numbers). As expected, the average mean proportion of noncrossover and crossover events carrying an uncorrected mismatch is significantly greater in *msh2* and *pms1* octads relative to wild-type (*P* < 0.0001; not shown on graph). No significant difference was found between the average mean proportion (**C**) or individual proportions (**D**) of crossovers versus noncrossovers carrying an uncorrected mismatch in wild-type octads, but in *ecm11* octads the proportion of crossovers carrying an uncorrected mismatch are significantly increased relative to the proportion of noncrossovers carrying an uncorrected mismatch. By contrast, the proportion of crossovers carrying an uncorrected mismatch are significantly decreased relative to the proportion of noncrossovers carrying an uncorrected mismatch in *msh2*, and in two out of three *pms1* octads. Table 3 lists the raw data per octad regarding uncorrected mismatches associated with the various categories of interhomolog recombination. Note chromosome 7 recombination events for two wild-type octads were excluded from these analyses, due to chromosome 7 disomy.

**Table 3.** Genome-wide errors in meiotic mismatch repair identified by Octad Rec-Seq. *(Relates to Figures 3, S2)* The table lists the raw number and percentages of interhomolog recombination events detected by Octad-RecSeq in four wild-type, five *ecm11*, two *msh2* and three *pms1* meioses. Uncorrected mismatches associated with interhomolog recombination events were detected by comparing the DNA sequences of mother versus daughter cells within octads using a custom R script (these companion DNA sequences, which represent the two products of a mitotic division, should be identical in the context of perfect meiotic mismatch repair). Data is plotted in Figures 3 and S2. Note chromosome 7 recombination events for two wild-type octads were excluded from these analyses, due to chromosome 7 disomy.

|  | Total<br>NCOs<br>(E1+E4) | Total COs<br>(E2+E3+E<br>5+E6+E7) | Total IH<br>events<br>(E1-E8) | Total IH<br>events w/<br>mismatch | % IH<br>events<br>(E1-E8) w/<br>mismatch | #NCO w/<br>mismatch | % NCO<br>(E1+E4) w/<br>mismatch | #CO w/<br>mismatch | % CO<br>(E2+E3+E<br>5+E6+E7)<br>w/<br>mismatch | Total<br>mismatches |  |  |  |  |  |  |
| --- | --- | --- | --- | --- | --- | --- | --- | --- | --- | --- | --- | --- | --- | --- | --- | --- |
| wt octad 1 no ch7 | 34 | 101 | 148 | 12 | 8.1 | 3 | 8.8 | 8 | 7.9 | 15 |  |  |  |  |  |  |
| wt octad 4 no ch7 | 28 | 81 | 125 | 15 | 12.0 | 3 | 10.7 | 10 | 12.3 | 20 |  |  |  |  |  |  |
| wt octad 7 | 33 | 114 | 167 | 20 | 12.0 | 4 | 12.1 | 14 | 12.3 | 23 |  |  |  |  |  |  |
| wt octad 8 | 42 | 127 | 190 | 16 | 8.4 | 4 | 9.5 | 12 | 9.4 | 18 |  |  |  |  |  |  |
| em11 octad 9 | 77 | 164 | 249 | 120 | 48.2 | 23 | 29.9 | 97 | 59.1 | 339 |  |  |  |  |  |  |
| ecm11 octad 10 | 107 | 233 | 355 | 189 | 53.2 | 39 | 36.4 | 147 | 63.1 | 537 |  |  |  |  |  |  |
| ecm11 octad 11 | 62 | 157 | 226 | 99 | 43.8 | 19 | 30.6 | 80 | 51.0 | 277 |  |  |  |  |  |  |
| ecm11 octad 12 | 95 | 162 | 270 | 97 | 35.9 | 26 | 27.4 | 68 | 42.0 | 177 |  |  |  |  |  |  |
| ecm11 octad 13 | 83 | 159 | 250 | 66 | 26.4 | 13 | 15.7 | 53 | 33.3 | 139 |  |  |  |  |  |  |
| msh2 octad 15 | 76 | 135 | 221 | 165 | 74.7 | 70 | 92.1 | 95 | 70.4 | 1199 |  |  |  |  |  |  |
| msh2 octad 16 | 72 | 135 | 215 | 170 | 79.1 | 69 | 95.8 | 101 | 74.8 | 1364 |  |  |  |  |  |  |
| pms1 octad 17 | 88 | 123 | 224 | 102 | 45.5 | 41 | 46.6 | 61 | 49.6 | 548 |  |  |  |  |  |  |
| pms1 octad 18 | 79 | 117 | 207 | 163 | 78.7 | 74 | 93.7 | 89 | 76.1 | 1111 |  |  |  |  |  |  |
| pms1 octad 19 | 109 | 116 | 238 | 194 | 81.5 | 105 | 96.3 | 89 | 76.7 | 1164 |  |  |  |  |  |  |
|  | #E1 w/<br>mismatch | %E1 w/<br>mismatch | #E2 w/<br>mismatch | %E2 w/<br>mismatch | #E3 w/<br>mismatch | %E3 w/<br>mismatch | #E4 w/<br>mismatch | %E4 w/<br>mismatch | #E5 w/<br>mismatch | %E5 w/<br>mismatch | #E6 w/<br>mismatch | %E6 w/<br>mismatch | #E7 w/<br>mismatch | %E7 w/<br>mismatch | #E8 w/<br>mismatch | %E8 w/<br>mismatch |
| wt octad 1 no ch7 | 3 | 8.8 | 6 | 8.2 | 1 | 10.0 | 0 | na | 1 | 10.0 | 0 | 0.0 | 0 | 0.0 | 1 | 7.7 |
| wt octad 4 no ch7 | 2 | 7.4 | 5 | 8.2 | 0 | 0.0 | 1 | 100.0 | 0 | 0.0 | 5 | 50.0 | 0 | na | 2 | 12.5 |
| wt octad 7 | 1 | 3.3 | 5 | 6.6 | 3 | 17.6 | 3 | 100.0 | 4 | 33.3 | 2 | 28.6 | 0 | 0.0 | 2 | 10.0 |
| wt octad 8 | 4 | 9.8 | 7 | 7.3 | 2 | 12.5 | 0 | 0.0 | 1 | 33.3 | 2 | 20.0 | 0 | 0.0 | 0 | 0.0 |
| em11 octad 9 | 19 | 27.5 | 40 | 47.1 | 26 | 66.7 | 4 | 50.0 | 8 | 88.9 | 19 | 76.0 | 4 | 66.7 | 0 | 0.0 |
| ecm11 octad 10 | 33 | 34.4 | 48 | 48.5 | 26 | 61.9 | 6 | 54.5 | 25 | 73.5 | 38 | 79.2 | 10 | 100.0 | 3 | 20.0 |
| ecm11 octad 11 | 13 | 24.1 | 34 | 40.0 | 13 | 59.1 | 6 | 75.0 | 13 | 76.5 | 16 | 55.2 | 4 | 100.0 | 0 | 0.0 |
| ecm11 octad 12 | 22 | 25.3 | 17 | 23.0 | 17 | 51.5 | 4 | 50.0 | 13 | 59.1 | 18 | 60.0 | 3 | 100.0 | 3 | 23.1 |
| ecm11 octad 13 | 12 | 15.8 | 20 | 22.5 | 8 | 25.8 | 1 | 14.3 | 11 | 73.3 | 13 | 56.5 | 1 | 100.0 | 0 | 0.0 |
| msh2 octad 15 | 52 | 89.7 | 32 | 47.1 | 35 | 89.7 | 18 | 100.0 | 19 | 100.0 | 7 | 100.0 | 2 | 100.0 | 0 | 0.0 |
| msh2 octad 16 | 60 | 95.2 | 39 | 56.5 | 36 | 90.0 | 9 | 100.0 | 12 | 100.0 | 9 | 100.0 | 5 | 100.0 | 0 | 0.0 |
| pms1 octad 17 | 29 | 43.9 | 18 | 33.3 | 21 | 48.8 | 12 | 54.5 | 4 | 80.0 | 18 | 85.7 | 0 | na | 0 | 0.0 |
| pms1 octad 18 | 60 | 93.8 | 40 | 62.5 | 35 | 89.7 | 14 | 93.3 | 2 | 100.0 | 9 | 100.0 | 3 | 100.0 | 0 | 0.0 |
| pms1 octad 19 | 84 | 96.6 | 46 | 69.7 | 29 | 87.9 | 21 | 95.5 | 9 | 100.0 | 4 | 66.7 | 1 | 50.0 | 0 | 0.0 |

Consistent with the observed increase in mismatch-associated recombination events, the total number of uncorrected mismatches detected for each Octad Rec-Seq dataset is elevated in *ecm11*, *msh2*, and *pms1* mutants relative to wild-type strains (Figure 3A light bars, Table 3). A total of 15, 20, 23 and 18 mismatches were detected among the four wild-type octad datasets. On the other hand, 339, 537, 277, 177, and 139 uncorrected mismatches were identified in the five *ecm11* datasets.

We next analyzed the number of uncorrected mismatches typically associated with poorly-repaired gene conversion tracts (tracts with at least one mismatch) in each strain. As the scatterplot in Figure 3B shows, *ecm11* mutants display a large number of recombination events harboring more than one mismatch, and several recombination events with more than five mismatches. However, the majority of recombination sites in *ecm11* mutants harbored fewer than five uncorrected mismatches; in fact the median number of uncorrected mismatches detected in all poorly-repaired recombination events for *ecm11* strains is two, while the median is one for wild-type, five for *msh2* and four for the *pms1* strain.

Taken together, our Octad-RecSeq data indicate that the SC central element protein, Ecm11, ensures robust mismatch repair at a large number of recombination sites genome-wide.

### Crossover-fated recombination intermediates are more susceptible than non-crossovers to errors in mismatch repair when Ecm11 is absent, but the reverse is true when Msh2 and Pms1 are absent

We used these Octad Rec-Seq data to investigate the relationship between mismatch repair errors and crossover outcome. First, we found that the percentage of recombination events carrying uncorrected mismatches in nearly every signature class is significantly increased in *ecm11*, *msh2,* and *pms1* octad datasets relative to wild type (Table 3, Figure S2). Thus, both majority and minority interhomolog recombination events are more likely to suffer from poor mismatch repair when Ecm11 is absent.

We next asked whether noncrossover and crossover events are equally likely to be susceptible to errors in mismatch repair. For wild type, the proportion of noncrossover events associated with an uncorrected mismatch is not significantly different from the proportion of crossovers associated with an uncorrected mismatch; this is true for the average values calculated from all four octads (Figure 3C) and for the values specific to each individual octad (Table 3, Figure 3D). However, each *ecm11* and *msh2* octad dataset, and two of the three *pms1* octad datasets, show a significant difference in the proportions of noncrossover versus crossover events carrying an uncorrected mismatch. Interestingly, in *ecm11* the fraction of crossover events carrying an uncorrected mismatch (∼50%) is significantly higher than the percentage of noncrossover events carrying an uncorrected mismatch (∼30%), while the reverse is true for *msh2* and *pms1* (Figure 3C, D, Table 3).

### The SC central element constrains gene conversion tract length, and crossover-associated conversion tracts with uncorrected mismatches average longer than their mismatch-free counterparts

Analysis of the octad datasets furthermore revealed a difference in gene conversion tract lengths in *ecm11* relative to wild type, particularly for crossover events (Figure 4A, B). The median gene conversion tract length for all crossover events detected in wild-type octads is 1793 nucleotides, while for *ecm11* the median crossover-associated tract length detected is 3352 nucleotides. *msh2* and *pms1* mutants show no significant deviation from wild type in crossover associated tract lengths, but show significantly shorter noncrossover tract lengths relative to wild-type (1100 and 1115 nucleotides, respectively, relative to the wild-type median of 1919 nucleotides). An increase in gene conversion tract length has been observed in other SC-deficient mutants, such as *msh4*, *zip3*, or strains homozygous for *spo11* hypomorphic alleles (Rockmill *et al*. 2013; Oke *et al*. 2014).

**Figure 4.**
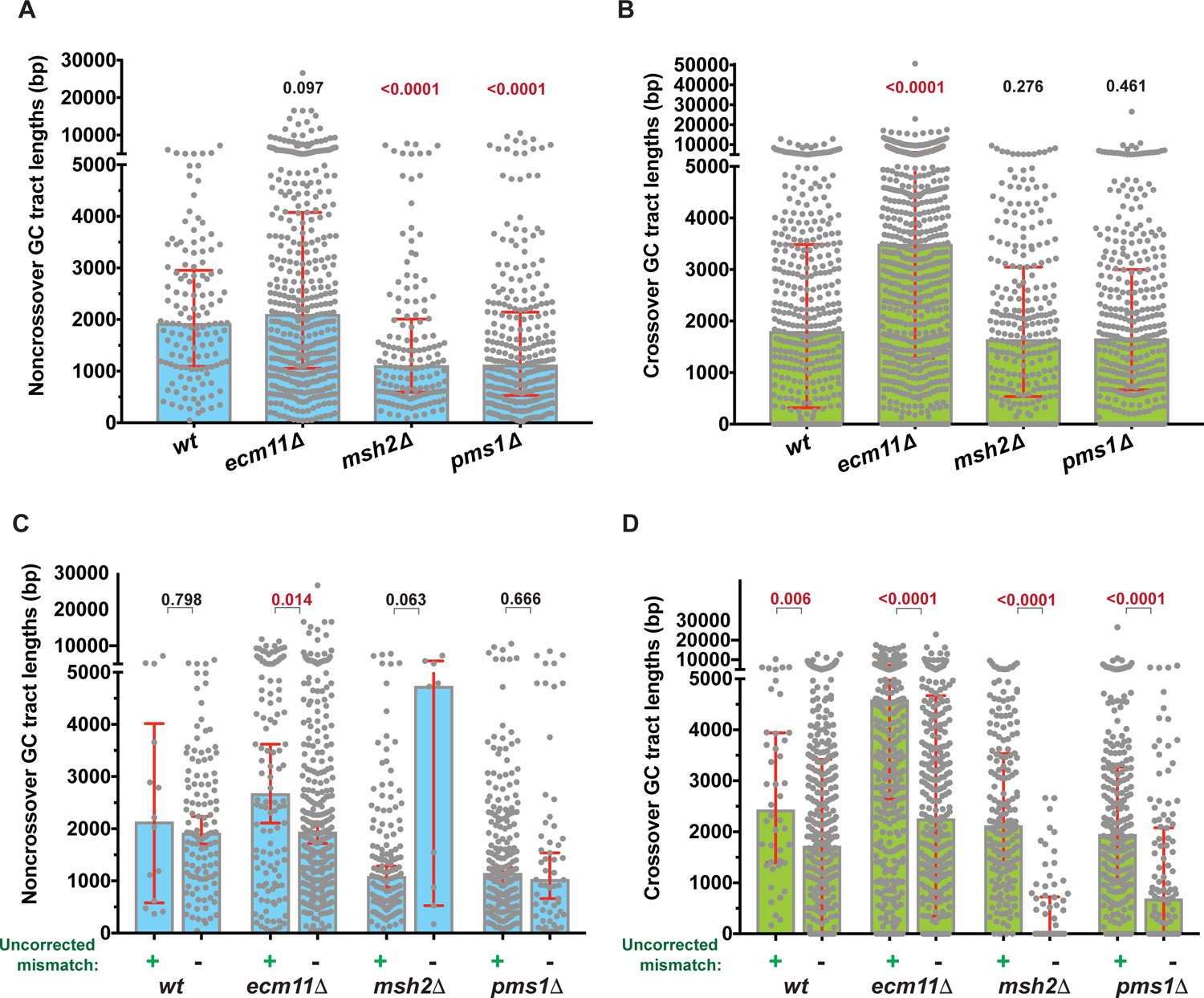
Crossover-associated gene conversion tract lengths are increased in *ecm11* mutants relative to wild-type, and crossover tracts carrying uncorrected mismatches average longer than their mismatch-free counterparts in all genotypes. Each circle on scatterplots in (**A**) and (**B)** represents a gene conversion tract length (# nucleotides) associated with a noncrossover (**A**) or a crossover (**B**) recombination event; data plotted derives from four wild-type, five *ecm11*, two *msh2*, and four *pms1* meioses (Octad Rec-Seq datasets; strains listed on the *x* axis). Height of the shaded area (blue in A, green in B) indicates a median value, while red error bars indicate interquartile range. Two-tailed *P* values (indicated above each column) using a Mann-Whitney test indicate a significant decrease in the lengths of noncrossover gene conversion tracts in *msh2* and *pms1* relative to wild type, and a significant increase in crossover gene conversion tract lengths in *ecm11* relative to wild type strains. Lower graphs plot noncrossover (**C**) or crossover (**D**) events carrying uncorrected mismatches (leftmost bars in each pair) separately from mismatch-free events (rightmost bars in each pair). Two-tailed *P* values using a Mann-Whitney test indicate no significant difference in gene conversion tract lengths between mismatch-containing and mismatch-free noncrossover recombination events in wild type, *msh2*, and *pms1* mutants, but mismatch-carrying noncrossover tracts are significantly longer than mismatch-free noncrossover tracts in *ecm11* mutants. Mismatch-carrying crossovers display significantly longer overall gene conversion tract lengths in all genotypes analyzed (*P* values indicated above each column). Note chromosome 7 recombination events for two wild-type octads were excluded from these analyses, due to chromosome 7 disomy. Additional data pertaining to the relationship between gene conversion tract length and frequency of mismatch repair errors are given in Figure 5, Figure S4, and Table S2, Table S3. Data regarding the relationship between recombination signature category and gene conversion tract length are given in Figure S3.

Strains bearing longer gene conversion tracts may display an increased number of uncorrected mismatches simply because their recombination intermediates carry a larger number of heterologous base pairs (SnPs) within the (longer) heteroduplex regions. If this is the case, we would expect recombination intermediates with longer gene conversion tracts to be more likely to bear uncorrected mismatches, relative to recombination intermediates with shorter gene conversion tracts. To ask this question we plotted mismatch-carrying gene conversion tract lengths separately from their mismatch-free counterparts within noncrossover and crossover classes for all genotypes (Figure 4 C, D). Analysis of noncrossovers revealed a small increase in the median tract lengths of mismatch-carrying events relative to mismatch-free events in wild type, *ecm11,* and *pms1* mutants, but a Mann-Whitney test determined the differences to only be significant in the case of *ecm11* (Figure 4C). On the other hand, the median tract lengths of crossovers carrying uncorrected mismatches was found to be substantially and significantly longer than that of mismatch-free crossover events in all four genotypes (Figure 4D). The positive correlation between tract length and mismatch repair errors in noncrossover and crossover recombination events in *ecm11* mutants is consistent with the possibility that an altered gene conversion tract length indirectly contributes to the elevated mismatch repair error phenotype observed in mutants missing the SC central element protein.

### Gene conversion tract length does not account for *ecm11*’s increased frequency of uncorrected mismatches

Is the increased frequency of mismatch-carrying recombination events in *ecm11* mutants *solely* due to the existence of abnormally long gene conversion tracts? Arguing against this idea is the fact that *ecm11* noncrossover gene conversion tract lengths are not significantly different from wild type, yet *ecm11* noncrossovers exhibit a higher frequency of uncorrected mismatches. We used two strategies to ask whether the increased frequency of mismatch-carrying crossovers observed in *ecm11* arises solely from an increased frequency of long crossover-associated gene conversion tracts. First, we classified recombination events into seven groups based on tract length size (arbitrary size ranges chosen were 0-1000bp; 1001-2000bp; 2001-3000bp; 3001-4000bp; 4001-10kbp; 10kbp-20kbp; 20kbp+), and calculated the frequency of events carrying an uncorrected mismatch in each group, for all four genotypes (Figure S4A, B, Table S2). If tract length were the only explanation for the difference between *ecm11* and wild type in frequency of recombination events with uncorrected mismatches, we expect *ecm11* to show a significant elevation in the proportion of mismatch carrying crossover events only in longer tract length categories. However, we found that *ecm11* octads display a significant increase in the proportion of crossover events with uncorrected mismatches in every tract length category (Figure S4B, Table S2).

We next addressed whether increased tract length alone accounts for elevated mismatch repair errors in *ecm11* mutants by calculating the frequency of uncorrected mismatches per nucleotide of gene conversion tract. If the increased errors in mismatch repair found in *ecm11* mutants is solely due to an overall increase in the cumulative length of gene conversion tracts, then the “per-nucleotide” frequency of uncorrected mismatches is expected to be similar between wild-type and *ecm11*. Instead, we found that the per-nucleotide frequency of mismatch repair errors is significantly (3-5 fold) higher in *ecm11* octad datasets relative to wild-type; this is true for both noncrossovers and crossovers (“All Noncrossovers” or “All Crossover” groups plotted in Figure 5A; “all events” columns in Table S3). Thus, the mismatch repair phenotype of *ecm11* mutants is not solely due to increased gene conversion tract length.

**Figure 5.**
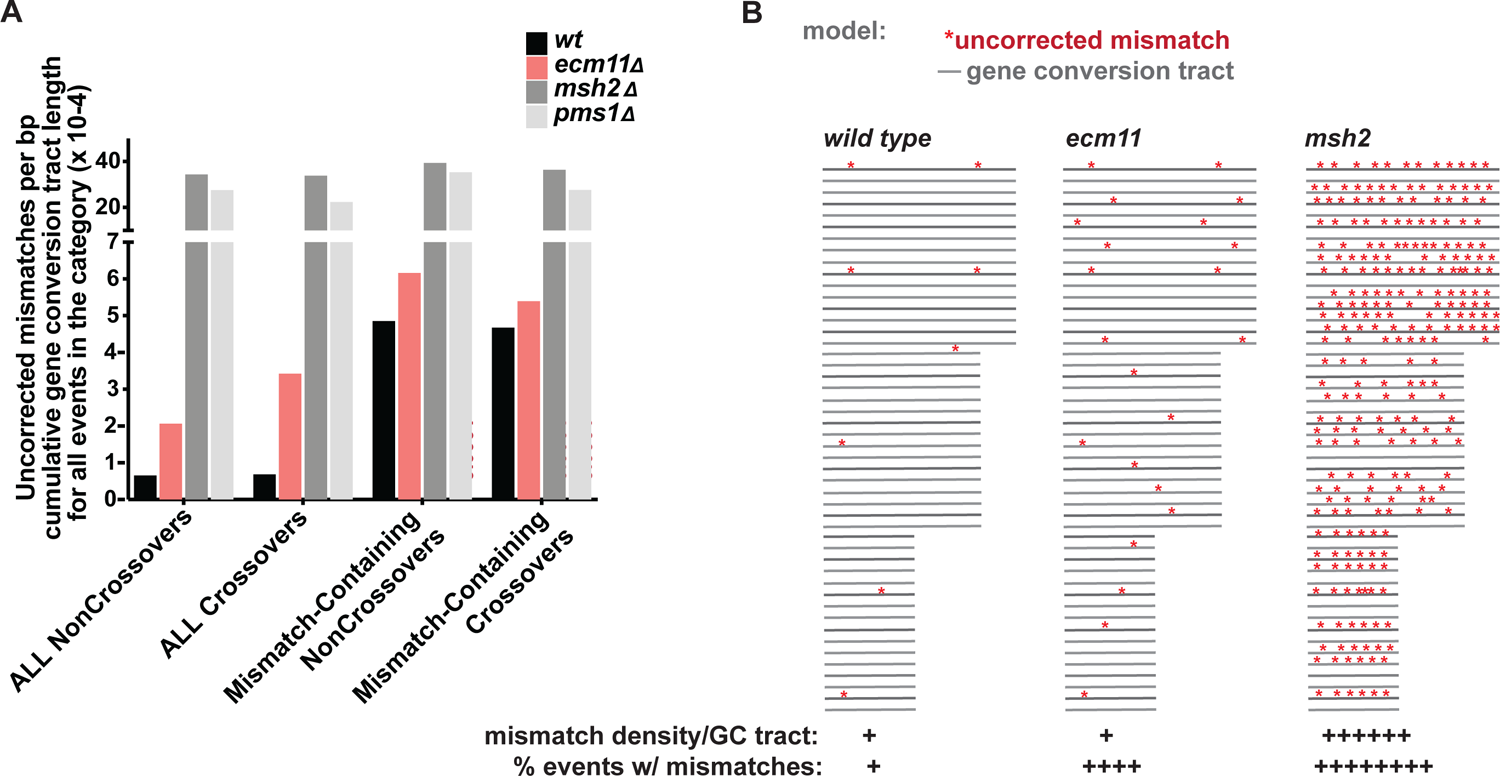
Increased gene conversion tract length fails to account for the elevated frequency of uncorrected mismatches in *ecm11* mutants. Graph in (**A)** plots the per-nucleotide frequency of uncorrected mismatches among noncrossover and crossover gene conversion tracts detected in four wild type, five *ecm11*, two *msh2*, and three *pms1* meioses (Octad Rec-Seq datasets). In the first group of four bars, an uncorrected mismatch per-nucleotide frequency for total *noncrossovers* is shown; this value was calculated by dividing the total number of uncorrected mismatches detected in a given genotype by the cumulative gene conversion tract length of all noncrossover events detected in that genotype. In the second group of bars, the mismatch per nucleotide frequency for total *crossovers* was calculated by dividing the total number of uncorrected mismatches detected in a given genotype by the cumulative gene conversion tract length of all crossover events detected in that genotype. The third and fourth groups of bars plots a per nucleotide mismatch frequency for *exclusively* uncorrected mismatch-carrying noncrossovers or crossovers (third and fourth groups respectively). In these latter calculations, the denominator used is the cumulative tract length of all uncorrected mismatch-carrying events detected in the genotype. Chi square with Yates Correction analysis of the “all event” (first two column groups) data indicates that mismatch repair errors arise significantly more frequently among both noncrossover and crossover events in *ecm11* relative to wild-type (thus the longer gene conversion tract lengths observed in *ecm11* mutants do not solely account for the frequency of uncorrected mismatches discovered in *ecm11* Octad Rec-Seq datasets). However, among exclusively those recombination events susceptible to inefficient mismatch repair, the per-nucleotide uncorrected mismatch frequency is not significantly different between *ecm11* and wild-type (*P* values calculated using Fisher’s Exact Test are indicated on the bars; red indicates statistical significance). The raw data plotted is given in Table S3. Note chromosome 7 recombination events for two wild-type octads were excluded from these analyses, due to chromosome 7 disomy. (**B)** Illustration compares the mismatch repair error phenotypes observed in wild-type*, ecm11*, and *msh2* strains. Gene conversion tracts of different lengths are indicated as grey lines; red asterisks correspond to uncorrected mismatch events. A majority of recombination events carry uncorrected mismatches in *msh2* mutants (Figure 3C), and the per base pair density of uncorrected mismatches within this category of gene conversion tracts is substantially higher than what is observed in wild type or *ecm11* mutants. In *ecm11* mutants, the density of uncorrected mismatches within gene conversion tracts is similar to wild type, but a greater proportion of recombination events belong to the “inefficient repair” class. This model is based on the two findings displayed in Figure 5A: that the frequency of uncorrected mismatches per base pair of total cumulative gene conversion tract length is significantly higher in *ecm11* mutants compared to wild type but the density of uncorrected mismatches among mismatch-containing gene conversion tract lengths is similar between the two genotypes. Together, the data indicate that wild-type strains exhibit a handful of recombination events with a low-level mismatch repair defect, and a larger proportion of recombination events are subject to this low-level mismatch repair defect when Ecm11 is absent. Note that increased gene conversion tract length and hyperrecombination phenotypes associated with *ecm11* are not depicted in this illustration.

Importantly, if we selectively consider only those gene conversion tract lengths associated with uncorrected mismatch-carrying events, the “per nucleotide” frequency of mismatch repair errors is similar between wild type and *ecm11* strains (“Mismatch-Containing” groups plotted in Figure 5A; “mismatch-carrying event” columns in Table S3). This latter result underscores a critical point, that the degree of mismatch repair deficiency is essentially the same among recombination events in an “inefficient MMR” class that exists in both wild type and *ecm11* mutants (illustrated in Figure 5B). Together our “per nucleotide” mismatch analyses raise the possibility that recombination intermediates susceptible to inefficient mismatch repair arise via the same mechanism in both *ecm11* and wild type strains: When Ecm11 (and the SC) is absent a large fraction of events are vulnerable to inefficient mismatch repair, whereas in wild type cells only a small number of events (perhaps those processed outside of the context of SC) fall into this inefficient repair class.

### Uncorrected mismatches in *ecm11* mutants are not likely due to excess DNA breaks or increased SnP density within gene conversion tracts

Ecm11 (or the SC) may act directly on recombination intermediates to ensure their efficient mismatch repair. We also considered the alternative possibility that mismatch errors arise in SC-deficient mutants because of a saturation of mismatch repair machinery. Prior studies indicate that budding yeast SC assembly prevents excess Spo11-initiated DSBs (Subramanian *et al*. 2016; Mu *et al*. 2020). To address the possibility that an excessive number of recombination events saturates mismatch repair machinery in SC-deficient mutants, we reduced DSBs using hypomorphic *spo11* alleles (Rockmill *et al*. 2013). We used *CRISPR*-Cas9-mediated gene targeting to create *zip1[Δ21-163]* strains homozygous for *spo11-179*, *spo11-32*, or *SPO11+*. We measured a reduction in chromosome III and VIII genetic map distances to wild-type levels or lower in *zip1[Δ21-163] spo11-179* and *zip1[Δ21-163] spo11-32* strains, while chromosome III and VIII genetic map distances displayed by our control strain, *zip1[Δ21-163] SPO11+*, were nearly identical to the original *zip1[Δ21-163]* strain (Figure S5A, Table S1). Despite the reduction in interhomolog recombination, post-meiotic segregation events involving *THR1* in our two *zip1[Δ21-163]* strains carrying *spo11* hypomorphic alleles occurred at an elevated frequency similar to the original *zip1[Δ21-163]* strain (Figure S5). Thus, the increase in mismatch-carrying recombination events observed in SC-deficient strains is not likely due to a saturation of the mismatch repair machinery due to excessive Spo11-mediated recombination initiation.

Mismatch repair machinery might also be saturated if SnP density is increased within gene conversion tracts. A gatekeeper activity has been attributed to mismatch repair proteins during meiosis, whereby nascent recombination intermediates involving some above-threshold degree of heterology are rejected via the activity of factors such as Msh2 (Borts *et al*. 2000). If the SC plays such a gatekeeper function, we expect gene conversion tracts to carry an increased SnP density in SC-deficient mutants relative to wild type. We therefore analyzed the SnP density across gene conversion tract lengths identified in our Octad Rec-Seq study. We found that SnP density values for both noncrossover and crossover gene conversion tracts in *ecm11* mutants do not significantly differ from wild type, which show, on average, 6.34 SnPs per nucleotide of noncrossover gene conversion tracts, and 4.8 SnPs per nucleotide of crossover gene conversion tracts (Figure S6; two-tailed *P* value comparing mutant to wild type is indicated above columns). We conclude that Ecm11 has no role in censoring heterology in recombination intermediates, and the increase in uncorrected mismatches observed in *ecm11* mutants is not due to an increase in SnP density within gene conversion tracts.

Interestingly, for both *msh2* and *pms1* strains, we found that the SnP density of noncrossover gene conversion tracts averaged two-fold higher than noncrossover gene conversion tracts found in wild type, and two-fold higher than *msh2* or *pms1* crossover gene conversion tracts (an average of 11.4 SnPs per nucleotide in *msh2* noncrossovers, versus 5.3 for *msh2* crossover events; 10.5 SnPs per nucleotide in *pms1* noncrossovers, versus 5.3 for *pms1* crossover events; Figure S6). The increased SnP density in noncrossover tracts is consistent with a role Msh2 and Pms1 proteins in rejecting strand invasion/transfer events that carry extensive sequence divergence (reviewed in Borts *et al*. 2000). However, the absence of a dramatically increased SnP density within crossover gene conversion tracts in *pms1* and *msh2* mutants suggests that these proteins do not play such a gatekeeper role, at least not to the same degree, for crossover-fated intermediates. These data thus suggest that Msh2 and Pms1 activities censor heteroduplex formation for noncrossover and crossover recombination intermediates in fundamentally different ways. We note that the lower density of SnPs within *pms1* and *msh2* crossover relative to noncrossover gene conversion tracts may be the reason that crossover-fated recombination intermediates in *msh2* and *pms1* mutants are less likely (relative to noncrossovers) to exhibit uncorrected mismatches (Figure 3C, D).

### SnPs near the termini of gene conversion tracts have an increased susceptibility to mismatch repair errors when Ecm11 is absent

In their pioneering analysis of genome-wide meiotic mismatch errors, (Mancera *et al*. 2011) revealed that uncorrected mismatches in wild-type meiotic recombination events are asymmetrically distributed toward the edges of gene conversion tracts. Using our Octad Rec-Seq datasets we calculated the frequency at which uncorrected mismatches are positioned at particular locations within gene conversion tracts of four wild type, five *ecm11*, and one *msh2* octad. We first assigned a standard relative position to every uncorrected mismatch: We defined the position of each uncorrected mismatch as a percentage of the distance from the midpoint (0%) and the closest terminus (100%) of its own gene conversion tract.

For example, an uncorrected mismatch positioned at the midpoint was assigned a position of 0%, a mismatch positioned half-way between the midpoint and either end of the gene conversion tract was assigned a position of 50%, and a mismatch positioned one nucleotide prior to either the start or end region of the gene conversion tract was assigned a position of 99%. Note that unlike other positions, position 100 is a bin that encompasses a variable number of nucleotides, due to the uncertainty in assigning gene conversion tract “start” and “end” points. A % distance value for every mismatch used in the analysis is plotted in Figure 6A, and the histogram in Figure 6B shows the frequency distribution of uncorrected mismatches at every position within gene conversion tracts.

**Figure 6.**
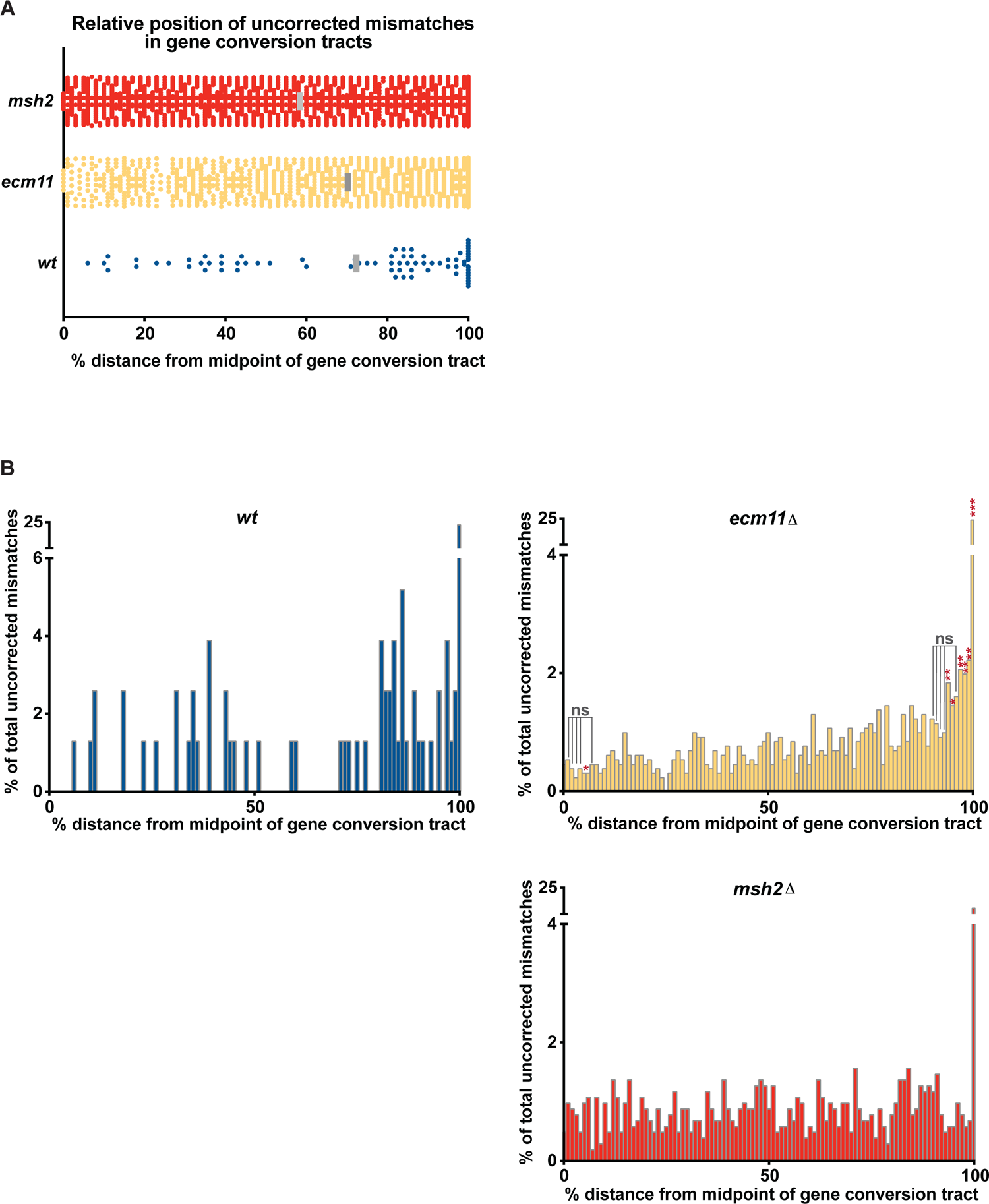
Uncorrected mismatches arising in wild type and *ecm11* mutant meiocytes display a slight positional bias toward the edges of gene conversion tracts. **A**) Plot gives the frequency of uncorrected mismatches positioned at particular locations within the gene conversion tracts of four wild type (blue circles), five *ecm11* (yellow circles), and one *msh2* (red circles) octad. A relative position was assigned to every uncorrected mismatch based on the distance between that position and the midpoint (0%) of its own gene conversion tract; a mismatch positioned half-way between the midpoint and either end of the gene conversion tract is assigned a position of 50%, and a mismatch positioned at the nucleotide just prior to the bin that defines either end of the gene conversion tract is assigned a position of 99%. Note that unlike the other positions, “position 100” is a bin encompassing a range of nucleotides that varies per event, due to the uncertainty in assigning gene conversion tract “start” and “end” points. The mean distance (grey bars) from the midpoint of a gene conversion tract is similarly larger in *ecm11* and wild type compared with *msh2* (suggesting a bias in mismatch position toward the ends of the gene conversion tract). Indeed, a Mann-Whitney test, which tests the null hypothesis that the distribution of values within each population is the same, found a significant difference between the distribution of uncorrected mismatch positions in *ecm11* and in wild type versus *msh2* (two-tailed *P =* < 0.0001, 0.0002, respectively), but no significant difference between the distribution of uncorrected mismatch positions in *ecm11* and wild type (two-tailed *P* = 0.639). The Mann-Whitney results thus suggest that the positions of uncorrected mismatches within *ecm11* and wild-type gene conversion tracts are indeed differently distributed from that of uncorrected mismatches within *msh2* gene conversion tracts. However, we note that a one-way analysis of variance (ANOVA) test and an unpaired t-test with Welch’s correction found no statistically significant difference between the means of the three populations (ANOVA *P*= 0.3691; t-test *P* > 0.999). This result underscores the fact thatwhile there may be a slight bias toward the ends, mismatches are left uncorrected at nearly any position within gene conversion tracts in all genotypes analyzed. (**B**) A difference in the distribution pattern of uncorrected mismatch positions within *ecm11* versus *msh2* gene conversion tracts is apparent in histograms that plot the frequency of uncorrected mismatches at every nucleotide position mapped from the midpoint of the gene conversion tract (0) to the edge (100%) for wild-type (blue), *ecm11* (yellow), and *msh2* (red). The histograms reveal a higher proportion of total uncorrected mismatches occurring at positions toward the edge of the gene conversion tracts in *ecm11* versus *msh2:* Red asterisks above bars in the *ecm11* graph indicate a significant difference relative to *msh2* in the proportion of uncorrected mismatches occurring at this position according to a Fisher’s Exact Test (*** for *P* <0.00001, ** *P* <0.01, * *P* <0.05). Grey “ns” indicators indicate no significant difference from *msh2* in frequency of uncorrected mismatches (*P*>.05) at this position. Bars with no markings were not evaluated for significance. (For all three strains, uncorrected mismatches appear most often at “position 100” of the gene conversion tract, but this may primarily be due to the fact that position 100 encompasses a bin of multiple nucleotides.)

Our analysis indicates that uncorrected mismatches occur across the entire gene conversion tract; we found no statistically significant difference between the means (grey bars in Figure 6A) of the populations of mismatch positions for wild type, *ecm11* or *msh2* (P= 0.3691). However, a Mann-Whitney Test found that positions of uncorrected mismatches within *ecm11* and wild-type gene conversion tracts are differently distributed from that of uncorrected mismatches within *msh2* gene conversion tracts (*P=*0.0002 for wild type versus *msh2* and < 0.0001 for *ecm11* versus *msh2* respectively; *P* = 0.639 for wild type versus *ecm11*). A difference in the distribution pattern of uncorrected mismatch positions within *ecm11* versus *msh2* gene conversion tracts is also apparent in histograms that plot the frequency of uncorrected mismatches at every nucleotide position (Figure 6B); these plots reveal a higher proportion of total uncorrected mismatches occurring at positions toward the edge of the gene conversion tracts in *ecm11* versus *msh2* mutants. Of the nine positions immediately adjacent to position 100, all nine show an elevated fraction of uncorrected mismatches in *ecm11* mutants relative to *msh2*, and for four positions the elevation is significant. Based on these data, we conclude that while SnPs positioned throughout gene conversion tracts in wild type and *ecm11* mutants are subject to inefficient mismatch repair, those near the extreme ends of the tract are particularly vulnerable.

### Deficient mismatch repair at *THR1* is more strongly linked to the absence of SC than it is to crossover pathway utilization

(Getz *et al*. 2008) built strains in which relatively high levels of post-meiotic segregation at *ARG4* and *HIS4* can be detected even in wild-type strains, owing to small palindromes that frequently escape mismatch correction near the DSB sites associated with these loci. This study showed that when MutSγ (Msh4) is removed, the proportion of mismatch-free crossover events is significantly reduced while the proportion of mismatch-associated crossovers (crossovers associated with a post-meiotic segregation event) remains equal or increased, thus the fraction of total recombination events with an uncorrected mismatch at each locus is increased in *msh4* mutants relative to wild type. This observation led the authors to suggest that interhomolog recombination intermediates processed independent of MutSγ are subject to poor mismatch repair relative to recombination intermediates processed via a MutSγ pathway. Consistent with this notion, we found that both *msh4* and *zip3* null mutants display a significant elevation in post-meiotic segregation at *THR1* (Figure 7, Table 4).

**Figure 7.**
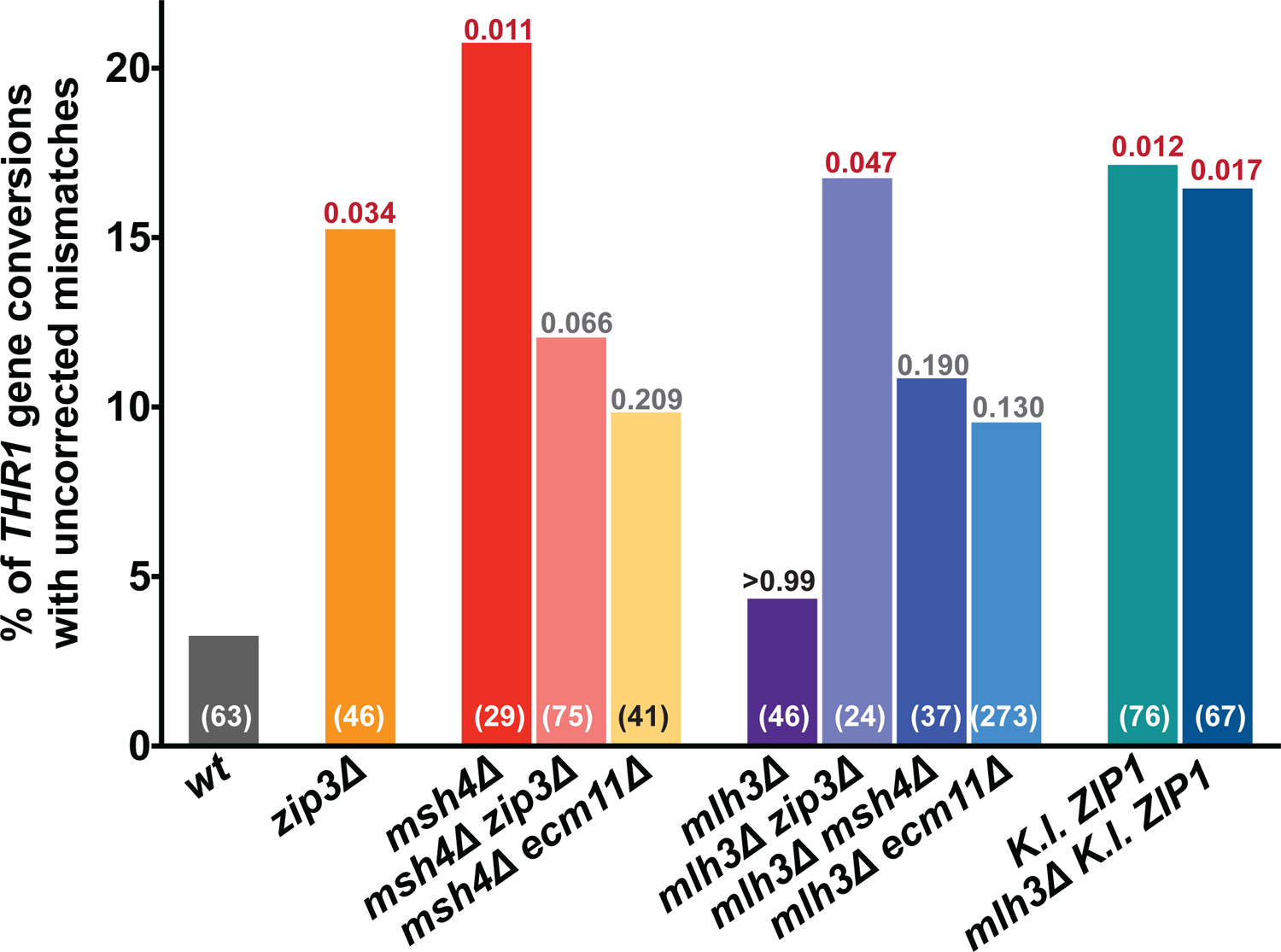
Some but not all MutSγ recombination-deficient mutants exhibit increased errors in mismatch repair at *THR1*. Bar graph plots the percentage of tetrads exhibiting non-mendelian segregation at *THR1* that display a phenotypically sectored colony on SC-Thr, reflecting the occurrence of an uncorrected DNA mismatch during meiosis. The total number of tetrads (meioses) observed with nonmendelian segregation at *THR1* is indicated (in white or black) at the bottom of the corresponding bar for each strain. Data plotted from mutants missing the pro-MutSγ recombination proteins, Zip3, Msh4 and Mlh3, as well as double mutants missing combinations of these proteins and/or SC structural proteins. In strains labeled *K.l. ZIP1*, the *S. cerevisiae ZIP1* gene ORF is replaced by the *ZIP1* gene from *Kluveromyces lactis; K.l.* Zip1 fails to support SC assembly in *S. cerevisiae* meiotic cells (Voelkel-Meiman *et al*. 2015). A two-tailed *P* value for each set of mutant data compared with wild type is indicated above individual columns (calculated using Fisher’s Exact Test; values in red are considered statistically significant, while those in grey not quite significant, and those in black not significant). See Table 4 for additional detail regarding the frequency of gene conversion at *THR1* and features of post-meiotic segregation events, and Table S1 for genetic map data associated with these strains.

**Table 4.**
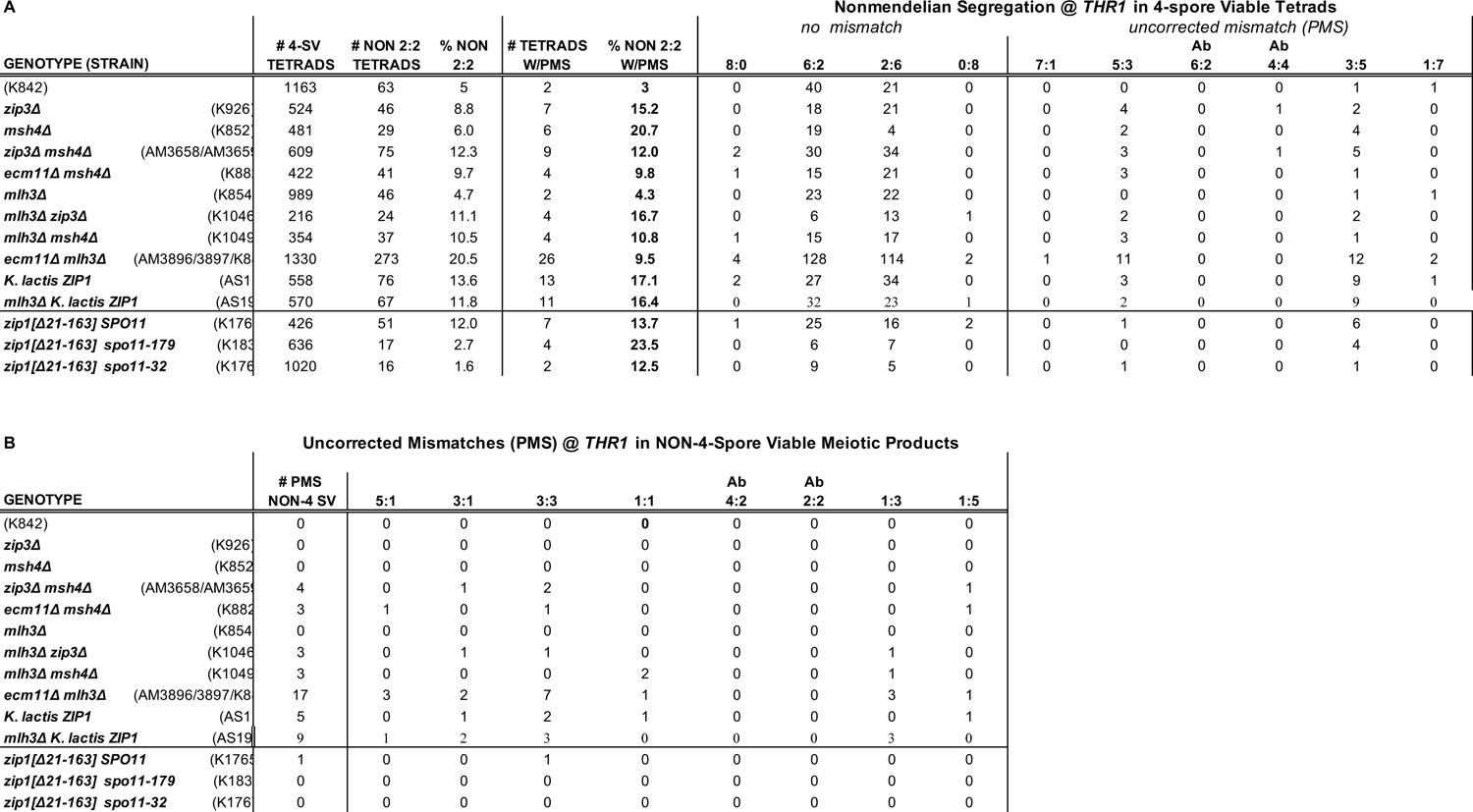
Errors in meiotic mismatch repair at THR1 are observed in mutants lacking some, but not all MutSγ-associated recombination pathway proteins. *(Relates to Figure 7)* Tables in (**A**) and (**B)** list meiotic segregation data for *THR1* in strains heterozygous for several genetic marker alleles, including *thr1-4*. **A**) Table gives the frequency of four-spore viable tetrads displaying gene conversion (non 2:2 segregation) at *THR1*, and the percentage of total tetrads associated with a failure in mismatch repair at this locus (tetrads that display a sectored colony on SC-Thr media), as well as the number of tetrads observed for each strain that fall into each category of non-mendelian segregation. **B**) The number of one-, two-, or three-spore viable meiotic products displaying a sectored colony on SC-Thr for each strain, and features of the post-meiotic segregation event (note ratios indicate whether the event occurred in a one-spore viable (1:1), a two-spore viable (3:1, Ab2:2, 1:3) or a three-spore viable (5:1, 3:3, Ab4:2, 1:5) meiosis. See Table S4 for strain names and full genotypes.

However, the possibility that mismatch repair is intrinsically more efficient for recombination intermediates handled by MutSγ is incompatible with the mismatch repair deficiency of *ecm11*, *gmc2,* and *zip1[Δ21-163]* mutants, where only MutSγ-dependent recombination appears to be elevated (Voelkel-Meiman *et al*. 2016). Furthermore, as in a previous study (Wang *et al*. 1999), we did not detect a significant increase in mismatch repair errors at *THR1* in strains lacking the MutLγ component Mlh3 (Figure 7, Table 4), which fail to form MutSγ crossovers (Argueso *et al*. 2004; Table S1). Among 989 tetrads from an *mlh3* diploid strain, only two out of 46 gene conversions at *THR1* (4.3%) were non-conventional, similar to the frequency of post meiotic segregation observed in our wild type strain. Taken together, these data show that an increase in mismatch repair errors (at least at *THR1*) is not correlated with (the presence or absence of) MutSγ-dependent crossing over.

SC assembly is compromised in strains missing certain MutSγ recombination factors such as Zip3 and Msh4 (Agarwal and Roeder 2000; Novak *et al*. 2001). If increased errors in mismatch repair observed in *msh4* and *ecm11* mutants are the result of absent or diminished SC, then we would expect SC assembly to be intact in *mlh3* mutants, which present no obvious defects in mismatch repair. To address this question, we surface-spread meiotic nuclei from wild type and *mlh3* meiotic cells onto glass slides, and evaluated the extent of SC assembly using antibodies against the transverse filament protein Zip1 and the central element protein Gmc2 (Figure S7). For this analysis we used strains lacking the transcription factor, Ndt80, which is required for progression beyond late meiotic prophase; *ndt80* mutant meiotic cells accumulate at a late meiotic prophase stage when SC structures are full-length (and joint molecules remain unresolved) (Allers and Lichten 2001a; Voelkel-Meiman *et al*. 2012). SC assembly was evaluated in diploid cells sporulated for 15 and 24 hours, as the majority of otherwise wild-type *ndt80* cell nuclei display completely full-length SC at the 24 hour time point (Voelkel-Meiman *et al*. 2012). We observed normal SC assembly in *mlh3* mutants: The SC structural proteins Zip1 and Gmc2 co-localize in linear structures to similar extents on surface-spread meiotic chromosomes from wild type and *mlh3* mutant meiocytes (Figure S7).

The presence of elevated post meiotic segregation at *THR1* in *zip3* and *msh4* but not *mlh3* mutants, in conjunction with the fact that SC assembly is unperturbed in *mlh3* mutants, leads us to favor the idea that Msh4 and Ecm11 guard recombination events from inefficient mismatch repair through their shared function in promoting SC assembly. This view is supported by our observation of elevated *THR1* post-meiotic segregation in *mlh3* double mutant strains that lack SC, such as the *mlh3 ecm11* double mutant, or an *mlh3* strain expressing *Kluveromyces lactis ZIP1*, which does not assemble SC (Voelkel-Meiman *et al*. 2015) (Figure 7; Table 4).

## Discussion

### SC central element proteins impact the interhomolog recombination process

While budding yeast SC is dispensable for interhomolog recombination *per se*, the data presented in this study point to several aspects of meiotic recombination that may be regulated by SC structure. Our analysis of post-meiotic segregation at *THR1* in various meiotic mutants reveals a strong correlation between robust SC assembly and efficient mismatch repair. Furthermore, high-throughput sequencing of octad meiotic products demonstrates that the SC structural protein, Ecm11, promotes efficient mismatch repair genome-wide at noncrossover and crossover recombination events. Our genome-wide sequence analysis moreover reveals additional impacts of Ecm11 and/or the SC structure on interhomolog recombination, including activities that 1) promote continuous gene conversion tracts, 2) discourage the use of more than one repair template, and 3) restrict gene conversion tract length. Interestingly, a similar lengthening of gene conversion tract length has been previously reported for SC-deficient strains, including *spo11* hypomorphic mutants (Rockmill *et al*. 2013; Oke *et al*. 2014).

Moreover, our prior observations raise the strong possibility that the SC structure can serve to influence whether MutSγ-associated double Holliday junction recombination intermediates are resolved in a biased or unbiased manner (illustrated in Figure 8A). *msh4* and *mlh3* single mutants display the same low level of crossovers when Ecm11 is present. However, *mlh3 ecm11* double mutants display a substantial increase in crossovers relative to *msh4 ecm11* double mutants, to a level that is precisely intermediate between the low crossover frequency of *msh4* or *mlh3* mutants and the elevated crossover frequency of *ecm11* single mutants (Voelkel-Meiman *et al*. 2016; relevant data also given in Figure S8). The elevated crossover frequency in *ecm11 mlh3* double mutants relative to *mlh3* single mutants suggests that removal of Ecm11 releases MutSγ-associated recombination intermediates from a constraint that ensures their resolution occurs in a biased manner. If both SC and MutLγ are present, MutSγ double Holliday junction structures resolve in a biased manner to become crossovers. If SC is present but MutLγ is missing (the *mlh3* mutant), MutSγ recombination intermediates still apparently resolve in a biased manner, but now toward the noncrossover outcome (Al-Sweel *et al*. 2017) using MutLγ-independent resolution enzymes. Finally, if the SC structure is absent alongside MutLγ (the *ecm11 mlh3* double mutant) MutSγ intermediates apparently resolve via an unbiased resolution pathway, giving 50% crossovers and 50% noncrossovers. We note that the increased crossover frequency in *ecm11 MLH3+* relative to *ecm11 mlh3* double mutants indicates that MutLγ itself can impose biased resolution toward a crossover outcome even if the SC is absent.

**Figure 8.**
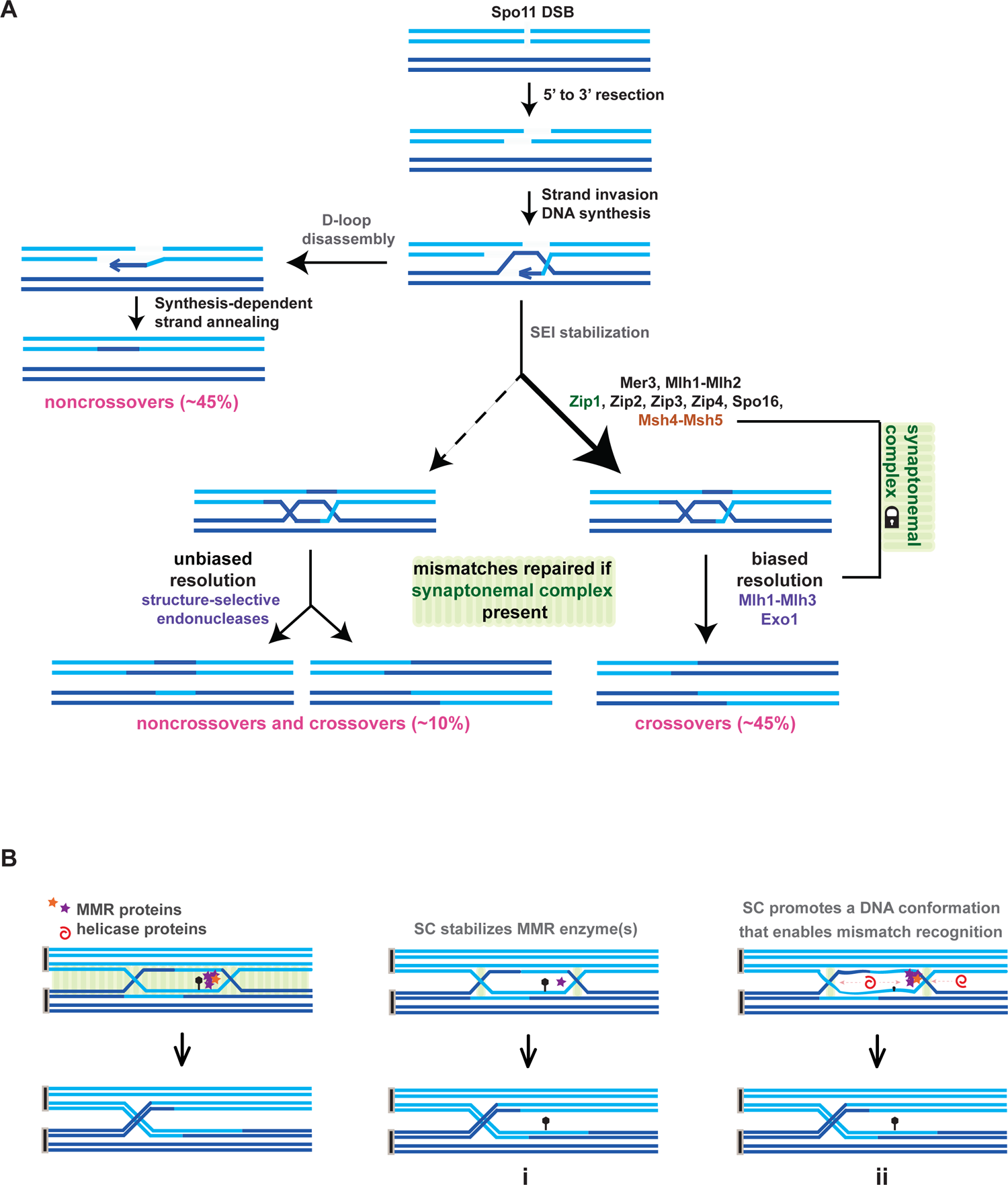
Roles for SC in processing interhomolog intermediates, and models for a possible mechanistic basis. Illustration in (**A**) is directly adapted from a meiotic recombination pathway illustration published in (Furman *et al*. 2021; Figure 1). Added to the model for meiotic recombination pathway choice is the apparent role for SCin constraining MutSγ (Msh4-Msh5) recombination intermediates to undergo biased resolution (this constrain is indicated by the lock icon), and the proposal that effective mismatch repair of joint molecule intermediates is dependent upon the presence of SC structure but not on which enzymes participate in resolution. In (**B**), a DNA joint molecule (double Holliday junction) is depicted in the physical context of tripartite SC (represented by green ovals spanning the distance between homologous chromosome axes). A mismatched base pair is indicated as a black lollipop within a heteroduplex region. After resolution of the joint molecule, the nonsister chromatids that had engaged in the recombination event are now structurally-continuous DNA duplexes free of mismatched bases, owing to mismatch repair enzyme activity (depicted as purple and orange stars). Images in “i” and “ii” illustrate possible molecular consequences of SC’s absence. The physio-chemical environment of the SC might serve to stabilize and thereby concentrate mismatch repair enzymes at the DNA joint molecule; when this function is missing some mismatches may be missed by the repair machinery (“i”). Alternatively, the protein architecture of the SC might directly or indirectly stabilize a double-stranded DNA conformation within heteroduplex regions of the joint molecule; when this function is missing, the structural properties of the DNA joint molecule may prevent some mismatches from being structurally accessible to the mismatch repair enzyme (“ii”). Structural proteins of the SC might influence joint molecule structure directly through a physical interaction with DNA, or indirectly by stabilizing another joint molecule-binding protein or by shielding DNA strands from enzymes that antagonize DNA duplex structure, such as helicases (depicted as red spirals).

Given MutSγ recombination complexes reside directly within or adjacent to SC structures (Voelkel-Meiman *et al*. 2019), a simple explanation for all of these findings is that the physical association of recombination intermediates with assembled or assembling SC structure influences the way such intermediates are processed. The SC structure might influence recombination intermediate processing through an activity that impacts recombination proteins themselves, or it could affect the way DNA structures are presented to recombination proteins. MutLγ, which appears to have a redundant function with SC in promoting a biased resolution outcome, is activated by polymer formation (Manhart *et al*. 2017). Thus one interesting possibility is that the polymer nature of the assembled SC and/or MutLγ physically bridges two junctions in a double Holliday junction structure in a way that directly facilitates their coordinated cleavage to give a biased resolution outcome.

### The basis for inefficient mismatch repair in *ecm11* and *msh4* mutants is likely the absence of SC

Here we present data that strongly links the synaptonemal complex structure to efficient mismatch repair. A prior study suggested that meiotic recombination events handled by the MutSγ pathway might be associated with a better mismatch repair outcome relative to MutSγ-independent recombination in budding yeast (Getz *et al*. 2008). Here the authors found that removal of the MutSγ component Msh4 leads to a diminishment in crossovers associated with conventional gene conversion relative to the level of crossovers associated with non-conventional conversion (post-meiotic segregation). Our observation of elevated post-meiotic segregation at *THR1* in *zip3* and *msh4* mutants is consistent with the possibility that a MutSγ recombination mechanism intrinsically associates with better mismatch repair relative to MutSγ-independent recombination. However, inconsistent with this possibility is the absence of an obvious mismatch repair defect in *mlh3* mutants (which also lack MutSγ-mediated crossovers), as well as the mismatch repair defect of *ecm11*, *gmc2,* and *zip1[Δ21-163]* strains, which exhibit increased MutSγ-dependent crossovers (Voelkel-Meiman *et al*. 2016). How are SC proteins that attenuate MutSγ recombination, as well as MutSγ itself (Msh4), involved in promoting efficient mismatch repair? We propose that the existence of SC structure determines whether interhomolog recombination intermediates are associated with effective mismatch repair in both *ecm11* and *msh4* mutants.

The SC appears to also constrain MutSγ joint molecule intermediates to a biased resolution pathway (Voelkel-Meiman *et al*. 2016), and biased resolution typically involves MutLγ. Thus, we also considered the possibility that - rather than mismatch repair being influenced by the SC itself-poor mismatch repair occurs when MutSγ recombination intermediates are resolved non-conventionally by a MutLγ-independent mechanism. However, the observation of mismatch repair errors in *msh4* mutants (which lack MutSγ recombination intermediates altogether) and the absence of mismatch repair errors in *mlh3* mutants (in which MutSγ intermediates are resolved by a MutLγ-independent mechanism) argue against this possibility (Getz *et al*. 2008; this work). We conclude that the role of both Ecm11 and Msh4 in mismatch repair is through promoting SC assembly, and that those recombination intermediates physically associated with the SC - regardless of whether they are handled by MutSγ - are susceptible to mismatch repair defects when SC is absent.

### Does the SC increase mismatch repair efficiency for a single kind of recombination intermediate?

Ligated joint molecules comprise the vast majority of recombination intermediates that exist when SC is full length in wild-type budding yeast, as demonstrated by physical gel analyses of recombination intermediates in cells blocked from progressing beyond a mid-meiotic prophase stage (Allers and Lichten 2001a; Sourirajan and Lichten 2008; de Muyt *et al*. 2012). Co-labeling of the SC and MutSγ-associated proteins on surface-spread meiotic chromosomes indicate that a large number of such joint molecule recombination intermediates are located at the midline of full-length SCs (Voelkel-Meiman *et al*. 2019), suggesting these recombination intermediate structures are directly peripheral to or embedded within SC structure. Joint molecules that form in a MutSγ-independent manner (responsible for the residual crossovers found in *msh4* mutants) may also embed within the SC structures that eventually form in such MutSγ-deficient strains. Thus, perhaps the SC promotes mismatch repair for *exclusively* ligated joint molecules.

Our Rec-Seq analysis indicates that crossover-fated recombination intermediates are more likely to carry uncorrected mismatches relative to noncrossovers when meiotic cells are missing Ecm11. This result is consistent with the idea that SC promotes efficient mismatch repair of joint molecule intermediates. The existence of noncrossovers with uncorrected mismatches in both *ecm11* and wild-type meioses raises the possibility that SC might also promote mismatch repair for intermediates that form through an SDSA mechanism. However, since SDSA-mediated noncrossovers form prior to bulk SC formation (Allers and Lichten 2001a) and independent of SC-promoting recombination proteins (Lynn *et al*. 2007; Borner *et al*. 2008), we favor a simpler explanation for the existence of noncrossovers with uncorrected mismatches in both *ecm11* and wild type strains - that they derive from a set of joint molecule intermediates processed by a MutLγ-independent mechanism. Although a vast majority of joint molecules become crossovers from biased resolution of Holliday junctions via a MutLγ-mediated mechanism in wild-type yeast (Argueso *et al*. 2004; Zakharyevich *et al*. 2012; Hunter 2015; Manhart and Alani 2016; Manhart *et al*. 2017), 10-20% of the joint molecules formed appear to rely on a (MutLγ-independent) structure-selective endonuclease pathway, involving Mus81-Mms4 and Yen1, which can generate crossovers and noncrossovers through unbiased cutting of DNA strands at each Holliday junction (de Los Santos *et al*. 2001; de Muyt *et al*. 2012; Zakharyevich *et al*. 2012; Figure 8). Noncrossover and crossover events carry uncorrected mismatches with equal frequency (∼10%) in wild-type hybrid strains (Mancera *et al*. 2011; this work). Thus, a tantalizing possibility is that uncorrected mismatches associated with wild-type recombination events arise from the 10-20% of joint molecules in the YJM796/S96 hybrid whose resolution is unbiased and mediated by the structure-selective endonuclease pathway. Perhaps these joint molecules are uniquely susceptible to mismatch repair errors because their resolution takes place outside of the context of the SC (discussed below).

### Uncorrected mismatches in *ecm11* and wild-type meiotic cells may arise via the same mechanism

Two key findings lead us to favor the view that uncorrected mismatches in *ecm11* and wild-type meiotic cells arise via the same mechanism. First, the distribution pattern of uncorrected mismatches in wild type and *ecm11* is slightly polarized toward the edges of gene conversion tracts, a pattern that is not observed for uncorrected mismatches in *msh2* mutants. Second, while the frequency of uncorrected mismatches per nucleotide of all gene conversion tracts is elevated in *ecm11* relative to wild type, the density of uncorrected mismatches within mismatch-containing gene conversion tracts is similar between the two genotypes (and much lower than the density of uncorrected mismatches in strains missing the mismatch repair enzymes Pms1 or Msh2). Based on these findings, we propose that 1) wild type and *ecm11* strains both harbor recombination events for which mismatch repair is similarly deficient, 2) the fraction of total recombination events that belong to this inefficient mismatch-repair class is significantly larger when SC central element proteins are absent. In our model, the few recombination intermediates susceptible to inefficient mismatch repair in wild-type strains are no different, at the molecular level, from recombination intermediates susceptible to inefficient mismatch repair in *ecm11* mutants, and perhaps arise (during otherwise wild-type meiosis) as a consequence of i) a local disruption in SC structure, ii) placement of a joint molecule outside of the context of SC, or iii) in the timing of a particular joint molecule’s resolution relative to SC assembly or disassembly.

### SC might impact mismatch correction by regulating the repair machinery or its substrate

The molecular mechanism underlying mismatch repair is best understood in prokaryotes: A MutS protein dimer recognizes and binds a mismatch in otherwise homoduplex DNA during replication, and a MutL dimer bridges the bound MutS with MutH, which nicks the newly replicated strand of DNA (through a strand discrimination mechanism involving DNA methylation). The nicked 3’ strand is resected until the misincorporated base is removed (Modrich 1996). Replicative mismatch repair in eukaryotes is carried out by a subset of MutS and MutL homolog family members (Kunkel and Erie 2015). During vegetative growth in budding yeast, Msh2, Mlh1, and Pms1 (Williamson *et al*. 1985; Reenan and Kolodner 1992; Prolla *et al*. 1994a), and to a lesser extent Msh3, Msh6, and Mlh3 (New *et al*. 1993; Marsischky *et al*. 1996; Flores-Rozas and Kolodner 1998) act as mismatch repair enzymes.

Homodimers of Msh2-Msh3 and Msh2-Msh6 bind mismatches *in vitro* as heterodimers (Alani 1996; Habraken *et al*. 1996), while Pms1 forms a heterodimer with Mlh1 (Prolla *et al*. 1994b). The mechanism that eukaryotes use to select which DNA to nick during DNA replication is not based on methylation but instead an orientation-specific interaction with the replication machinery (Kadyrov *et al*. 2006; Pillon *et al*. 2010; Pluciennik *et al*. 2010; Pillon *et al*. 2015).

During meiotic recombination, mismatch repair of heteroduplex determines whether a DNA duplex will carry original sequence or be converted to the sequence of its homologous partner. Meiotic mismatch repair in yeast relies on Msh2, Mlh1 and Pms1 (reviewed in Borts *et al*. 2000). However little is known about the molecular mechanism(s) of meiotic mismatch repair, including at what stage(s) mismatches are detected and repaired (i.e. during strand invasion, D-loop formation, strand extension by polymerase, or after ligated joint molecules have formed), or how the direction of repair is determined toward conversion or restoration within heteroduplex (i.e. which strand is nicked to initiate repair). Analysis of uncorrected meiotic mismatches in *pms1* and *msh2* single and double mutants indicate that the two enzymes appear to at least in part act in parallel, independent pathways (Alani *et al*. 1994; Borts *et al*. 2000; Hoffmann and Borts 2004), but distinguishing molecular features of these pathways remain unclear.

How might SC impact mismatch repair mechanism(s)? The SC does not play a central role in the enzymology of mismatch repair; this is clear from the dramatic difference in total number of uncorrected mismatches between *ecm11* and *msh2* or *pms1* mutants. Our data also argues against the possibility that mismatch errors in SC-deficient mutants arise indirectly through a saturation of mismatch repair machinery due to excessive recombination or elevated SnP density.

As discussed above, previously published genetic data suggests that assembled SC appears to couple MutSγ recombination intermediates to a biased resolution pathway (Voelkel-Meiman *et al*. 2016; Figure 8A); perhaps the SC promotes efficient mismatch repair through an associated mechanism. For example, a molecular aspect of the biased resolution mechanism might stabilize mismatch repair protein(s) at recombination sites. We note that the absence of a mismatch repair defect in *mlh3* strains indicates that such a hypothetical feature of the resolution mechanism is not the MutLγ complex itself, nor is it dependent on MutLγ. Alternatively, the physio-chemical properties of structural proteins that create the SC “compartment” at the recombination site might serve to concentrate or trap mismatch repair enzymes at the DNA joint molecule (Figure 8B, “i”). The central region of the SC (comprised of transverse filament and central element) is assembled by proteins that, while maintaining a specific orientation within the overall structure, interact with one another via a range of binding affinities, and the capacity for SC central region to be dissolved by aliphatic alcohols indicates that SC architecture relies heavily on a large number of weak hydrophobic interactions (Rog *et al*. 2017). Furthermore, SC central region proteins exhibit dynamic behavior: in budding yeast both transverse filament and central element proteins continuously incorporate into preexisting full-length SCs (Voelkel-Meiman *et al*. 2012) and in other organisms protein subunits of the SC have the capacity to rearrange within the structure (Nadarajan *et al*. 2017; Pattabiraman *et al*. 2017; Rog *et al*. 2017). These observations indicate that structural subunits may not necessarily impose a barrier to the movement and/or rearrangement of recombination proteins, such as MutSγ or MutLγ complexes, within the SC. Instead, the unique chemical and structural properties of an enveloping SC may serve a chaperone-like role in regulating recombination protein interaction or aggregation behavior by stabilizing interactions between such proteins or by constraining their movement to a limited terrain (i.e. along the length of the chromosome axis). In light of this idea it is interesting that MutLγ is activated by polymer formation (Manhart *et al*. 2017).

Another possibility is that the SC acts in a chaperone-like manner to promote a certain structural conformation of the DNA joint molecule itself (Figure 8, “ii”). The MutS family of mismatch repair enzymes binds homoduplex DNA to “scan”, then recognizes/binds a mismatch with higher affinity owing to the ability of that nucleotide to distort, when bound, within the otherwise uniform homoduplex DNA landscape (Lamers *et al*. 2000). Numerous weak interactions between the SC proteins and DNA strands could theoretically serve to stabilize a DNA homoduplex conformation within heteroduplex regions of the joint molecule, allowing repair enzymes to remain bound to the DNA in “scan” mode or allowing mismatched nucleotides to manifest structurally in a way that enables high affinity binding by repair enzymes. Such a role for the SC in stabilizing a homoduplex conformation of DNA might be particularly important toward the edges of the gene conversion tract where criss-crossed single strands of DNA (Holliday junctions) exist; such junctions may have an intrinsic susceptibility to losing homoduplex conformation. This idea is interesting in light of the polarized distribution of uncorrected mismatches toward the edges of gene conversion tracts in *ecm11* mutants.

The SC could alternatively influence the structural conformation of heteroduplex regions indirectly, by stabilizing one or more recombination proteins that bind to the joint molecule structure or by shielding the recombination intermediate from enzymes that antagonize DNA duplex structure, such as helicases. This last idea is interesting in light of the finding that the SC-promoting recombination proteins, including Zip1 and MutSγ, protect interhomolog recombination intermediates from dissolution by the Sgs1 helicase (Jessop *et al*. 2006). A role in regulating the structural conformation of a DNA joint molecule and/or protecting it from being accessible to DNA-targeted enzymes could also underlie how the SC normally prevents abnormally long gene conversion tracts (Rockmill *et al*. 2013; Oke *et al*. 2014; this work) and perhaps how the SC structure constrains MutSγ intermediates to undergo biased resolution.

## Methods

### Strains

Strains (Table S4) are of the BR1919-8B background (Rockmill and Roeder 1998), except for those listed under the heading “Octad Rec Seq Experiments”, which are derived from a cross between YJM789 and S96 genetic backgrounds (Wei *et al*. 2007; Anderson *et al*. 2011). Knockout mutants were created by standard recombination-based gene targeting procedures or CRISPR-Cas9 mediated allele replacement as described in (Voelkel-Meiman *et al*. 2019). Diploid strains used in Octad Rec-Seq experiments were created by crossing a YJM789 parent with an S96 parent. The YJM789 isolate used in these experiments carries an allele of the *AMN1* gene *[A1103T]* (Mancera *et al*. 2011) which helps facilitate the mechanical separation of mother and daughter cell products of the first vegetative cell division.

### *THR1* PMS identification and crossover analysis

Post meiotic segregation at the *THR1* locus is detectable as sectored growth of a spore colony on SC-Thr media. Such sectored colonies were additionally verified by isolating cells from either side of the sector boundary within the corresponding colony on rich media, and checking these isolates for differential growth on SC-Thr. Conventional segregation of several other heterozygous markers ruled out the possibility that sectored colonies arise from accidental placement of spores directly adjacent to one another during the dissection process; the colonies that grew in a sectored manner on SC-Thr always presented a non-sectored growth phenotype on other selective media (i.e. Hyg, Nat, SC-Ura, SC-Trp, SC-Ade, SC-Lys, SC-His media).

Genetic crossover data was processed using an Excel Linkage Macro program created by Jonathan Greene (Rhona Borts, pers. comm) and donated by Eva Hoffmann (University of Copenhagen, Denmark). Crossover values and standard errors were obtained usimg the Stahl lab online tools (https://elizabethhousworth.com/StahlLabOnlineTools/), with the method of Perkins (Perkins 1949).

### Canavanine resistance mutator assay

Mutator assay was conducted as in (Argueso *et al*. 2003). Eleven single colonies from wild type, *ecm11*, or *pms1* diploids were patched onto media containing 60μg/mL canavanine, and incubated for 3-4 days at 30°C. The median number of papillae from each of the eleven patches was recorded for each genotype. The average of median values from four repetitions are plotted in Figure 1D.

### Octad isolation and preparation for Illumina Sequencing

Tetrads from control, *ecm11*, *msh2*, *pms1* hybrid strains (Table S4) were dissected onto rich media, and individual spores were then continuously monitored for the onset of cell division. Once mitosis/cytokinesis had progressed to the extent that mother and daughter cells could be manually separated (sometimes several hours after spore placement onto rich media), the glass dissection needle was used to separate mother from daughter on the plate. Spore viabilities of hybrid strains is given in the table below:

| Genotype (strain) | Tetrads dissected | 4 spore viable | 3 spore viable | 2 spore viable | 1 spore viable | 0 spore viable | Spore viability (%) |
| --- | --- | --- | --- | --- | --- | --- | --- |
| wild type (AM3740) | 66 | 40 | 15 | 6 | 3 | 2 | 83 |
| *ecm11* (AM3735) | 155 | 34 | 60 | 36 | 18 | 7 | 65 |
| *msh2*<br>(AM3736) | 102 | 16 | 16 | 30 | 29 | 11 | 49 |
| *pms1*<br>(AM3738) | 92 | 17 | 18 | 28 | 21 | 8 | 54 |

Before preparing genomic DNA, each tetrad used in our analysis was first verified by observing 2:2 segregation of six to eight distinct heterozygous markers. This was done using PCR with custom primers designed to discriminate between S96 and YJM789 genome sequence. Eight cells comprising the four mother and four daughter cells of 4 wild-type, 5 *ecm11*, 2 *msh2* and 3 *pms1* octads were individually cultured for genomic DNA isolation followed by bar-coded library preparation, which was performed as in (Oke *et al*. 2014).

### Data Analysis

The ReCombine software package (Anderson *et al*. 2011; Oke *et al*. 2014) was used on Fastq files from Illumina’s Casava pipeline to align all sequences to reference genomes, genotype markers, and designate CO and NCO locations for each mother or daughter “tetrad” component of the octad. Recombination events were compared between each pair of mother and daughter spores and mismatches were identified using a custom R script. Indels were removed. The data will be deposited in Dryad once a manuscript submission number is available. Statistical tests were performed using GraphPad Prism software (www.graphpad.com).

### Cytology

Meiotic cell nuclei were surface-spread to glass slides and imaged as described in (Voelkel-Meiman *et al*. 2016). Affinity purified rabbit anti-Zip1 and mouse anti-Gmc2 (Voelkel-Meiman *et al*. 2019) were used at 1:100 and 1:800 dilutions, respectively. Alexa Fluor-conjugated anti-rabbit and anti-mouse secondary antibodies purchased from Jackson Immunoresearch were used at a 1:200 dilution. Slides were mounted in a glycerol-based media containing DAPI. Chromosomes were imaged on a Deltavision deconvolution microscope (General Electric) adapted to an Olympus (IX71) microscope. Images shown are projections of two to three individual focal planes along the z axis.

## Acknowledgements

We thank Drs. Beth Rockmill and Tim Nelson, their dog Ecco, as well as all members of the Fung lab, for their hospitality during the week spent preparing samples for Illumina sequencing.

---

**Figure S1.**
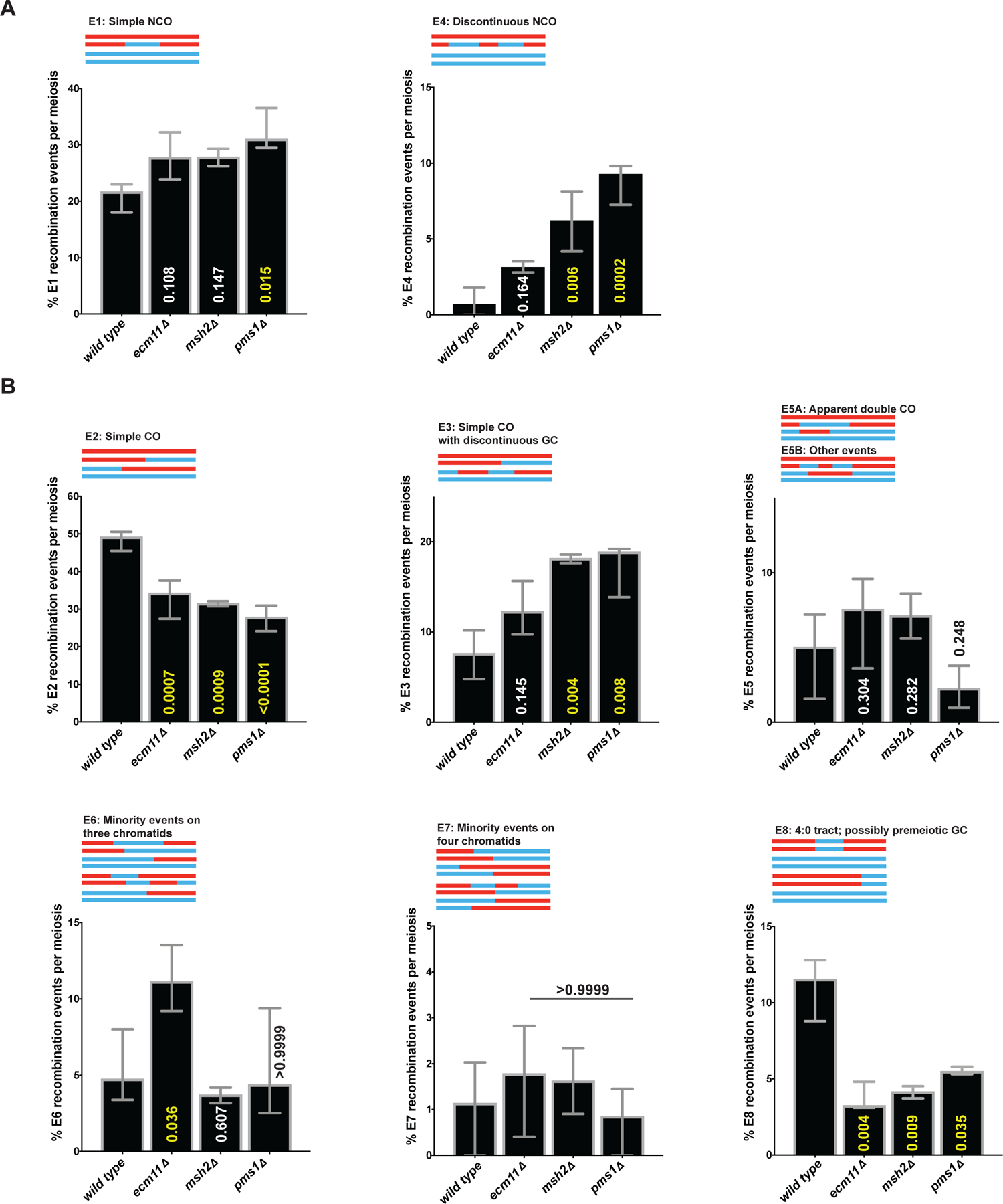
Genome-wide sequence analysis of *ecm11* meiotic products indicates a role for SC central element in limiting both majority and minority interhomolog recombination events. ***(Relates to Figure 2***) Bar graphs indicate the proportion of specific interhomolog recombination events found in four wild type, five *ecm11*, two *msh2*, and three *pms1* Octad Rec-Seq datasets; median and range values for each strain are plotted. *P* values obtained by Fisher’s Exact Test (indicated in yellow if significant, white if not significant) indicate whether the average proportion of a particular class of recombination events in each mutant is significantly different from the average proportion found in wild type. Recombination event categories are described in Figure 2 legend, and data plotted in these graphs is listed in Table 2. Note chromosome 7 recombination events were excluded for all octads, due to chromosome 7 disomy in two wild-type samples.

**Figure S2.**
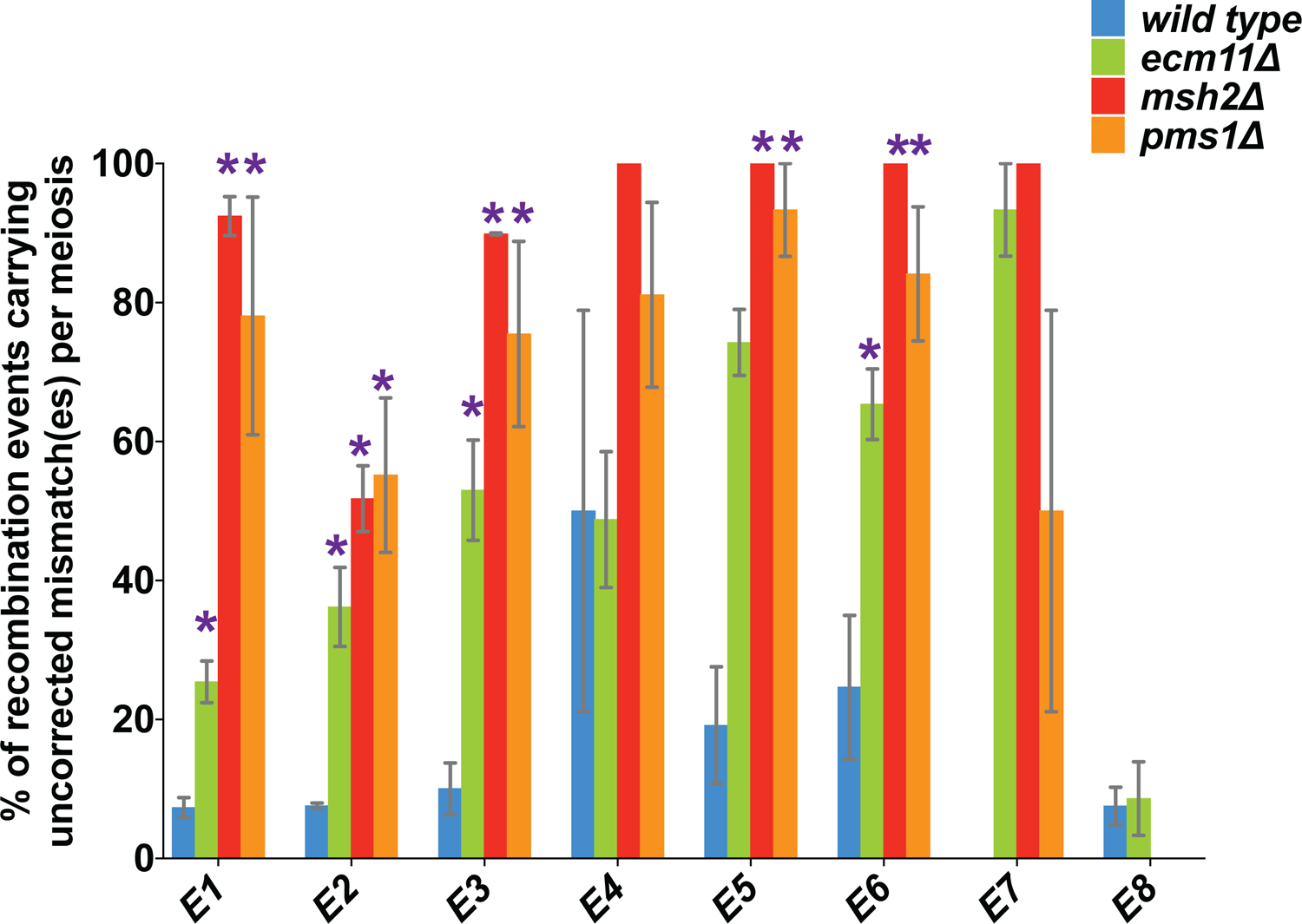
*ecm11* mutants exhibit an increased number uncorrected mismatch-carrying events in several categories of recombination. *(Relates to Figure 3)* Column graph plots the median and range percentage of specific classes of recombination events (indicated on the *x* axis) that display uncorrected mismatches in four wild-type (blue), five *ecm11* (green), two *msh2* (red), and three *pms1* (orange) octads. For each class of recombination event, strains showing significant differences relative to wild type are labeled with a purple asterisk (two-tailed *P* values obtained using a Fisher’s Exact Test). The proportion of E1 recombination events with uncorrected mismatches for *ecm11*, *msh2,* and *pms1* are significantly increased relative to wild type (*P*=0.045 <0.0001, <0.0001 respectively). The proportion of E2 and E3 recombination events with uncorrected mismatches is also significantly increased in *ecm11*, *msh2,* and *pms1* (*P* <0.0001 for E2 in each strain; *P=* 0.04, <0.0001, 0.0007 respectively, for E3). The proportion of E5 recombination events with uncorrected mismatches is not significantly different between wild-type and *ecm11* (*P*=0.69), but is significantly different between wild-type and *msh2,* or *pms1* (*P=*0.0006, and 0.028 respectively). Finally, the proportion of E6 recombination events with uncorrected mismatches is significantly different from wild-type in *ecm11*, *msh2*, and *pms1* (*P=*0.045, 0.007, 0.019, respectively). Note chromosome 7 recombination events for two wild-type octads were excluded from the analysis due to chromosome 7 disomy.

**Figure S3.**
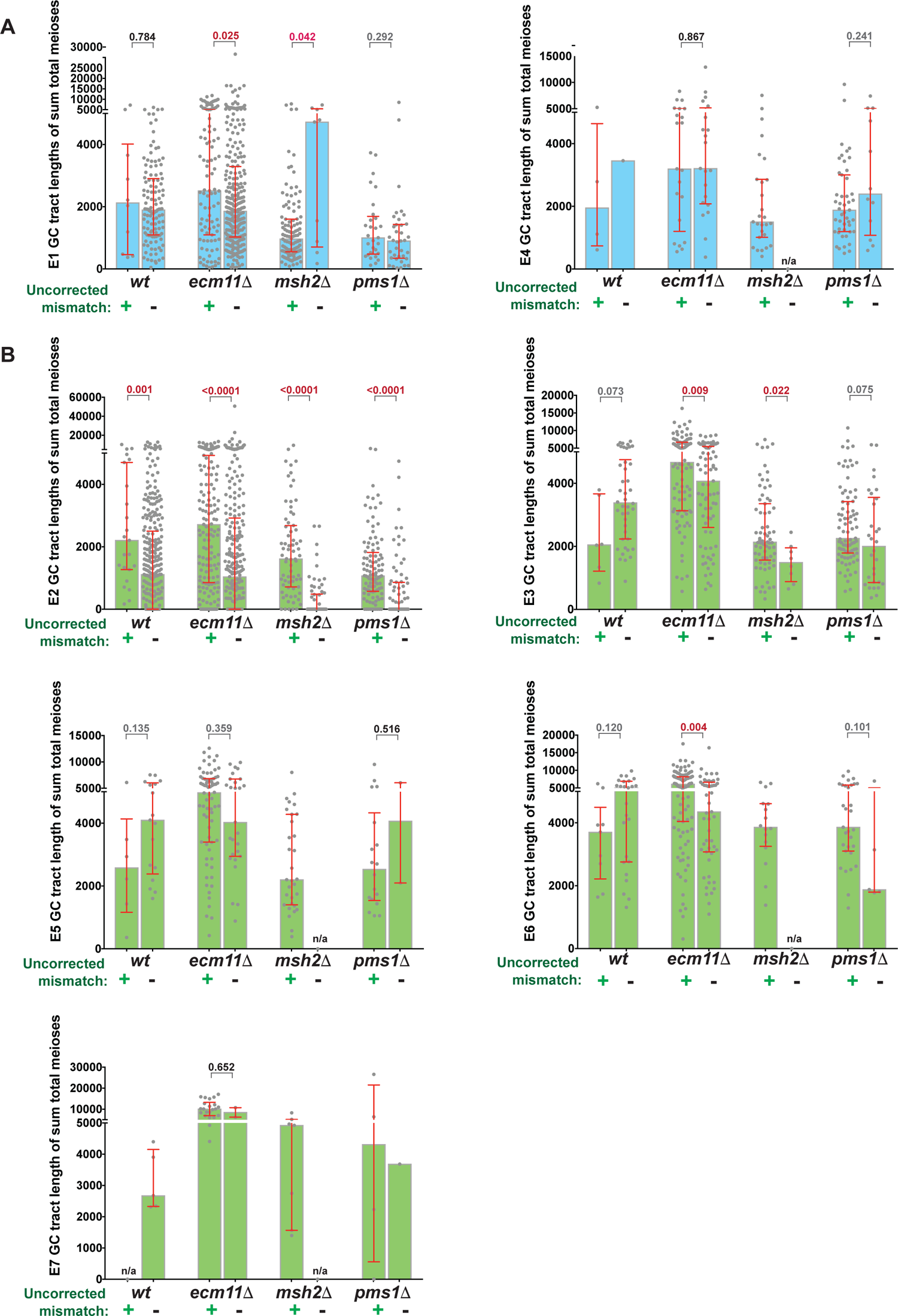
Gene conversion tracts with uncorrected mismatches often average longer than their mismatch-free counterparts in *ecm11* mutants. *(Relates to Figure 4)* Each circle on the scatterplots represents a gene conversion tract length (# nucleotides) associated with a noncrossover (light blue shading, **A**) or crossover (green shading, **B**) class of recombination events. Total gene conversion tract lengths with each type of recombination signature (E1-E7) from four wild-type, five *ecm11*, two *msh2*, and four *pms1* meioses (Octad Rec-Seq datasets) are plotted in separate columns depending on whether the event carries an uncorrected mismatch (indicated below with a + or -; strains listed on the *x* axis). Height of the shaded area indicates a median value for each group, while red error bars indicate interquartile range. Two-tailed *P* values using a Mann-Whitney test are indicated above the error bars (red if significant). Note chromosome 7 recombination events for two wild-type octads were excluded from these analyses, due to chromosome 7 disomy.

**Figure S4.**
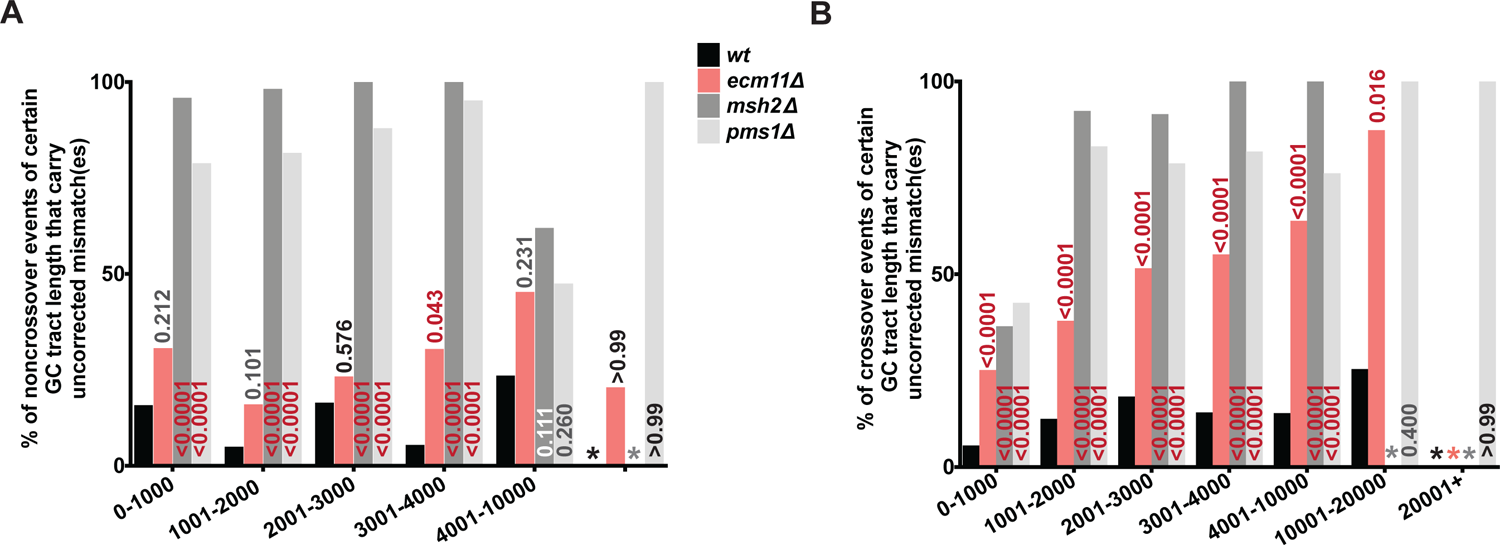
Elevated frequency of uncorrected mismatches in *ecm11* mutants regardless of gene conversion tract length. *(Relates to Figure 5,* Table S2) Column graphs plot the percentage of uncorrected mismatch-carrying noncrossover (**A**) or crossover (**B**) events with gene conversion tracts of distinct length ranges (indicated on the *x* axis). Data plotted is consolidated from all genome-wide recombination events identified in four wild-type (black bars), five *ecm11* (pink bars), two *msh2* (gray bars) and three *pms1* (silver bars) meioses (Octad Rec-Seq datasets). For each category of tract length, a Fisher’s Exact Test was used to determine whether the proportion of uncorrected mismatch-carrying recombination events is significantly different in any of the mutant strains relative to wild-type (*P* values are indicated on the graph; red indicates statistical significance). The proportion of noncrossovers carrying an uncorrected mismatch in *ecm11* is only significantly greater than wild-type in the 3001-4000 tract length category, but the proportion of crossovers carrying an uncorrected mismatch in *ecm11* is significantly greater than wild-type for every tract length category. The raw data plotted in (**A**, **B**) is given in Table S2. Note chromosome 7 recombination events for two wild-type octads were excluded from these analyses, due to chromosome 7 disomy.

**Figure S5.**
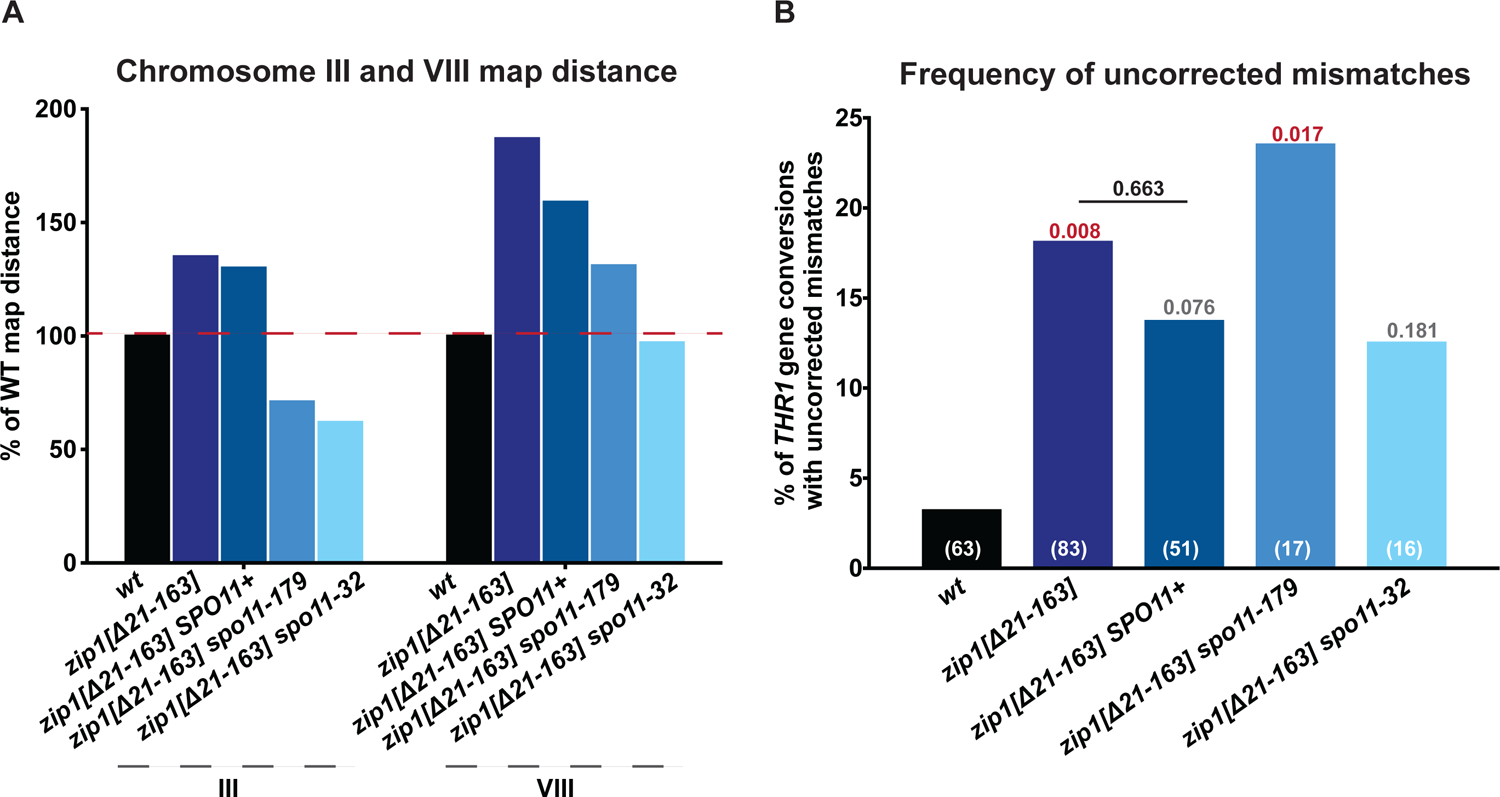
The mismatch repair defect of SC-deficient mutants is not likely due to saturation of mismatch repair machinery caused by elevated DNA double strand breaks. Graph in (**A)** plots the genetic map length of chromosome III (left cluster) and chromosome VIII (right cluster), in *zip1[Δ21-163]* diploids homozygous for either *SPO11+* (navy bar) or two different *spo11* hypomorphic alleles (light blue bars). Five genetic markers spanning chromosome III and four genetic markers, including *THR1/thr1-4*, along the length of chromosome VIII were used to calculate map distances (See Table S1 for full genetic map data). Summed genetic map distances between all intervals on III and VIII are displayed as a percentage of the wild-type value. A reduction in the genetic map length relative to both *zip1[Δ21-163] SPO11+* and wild type is observed in each strain carrying a *spo11* hypomorphic allele. Graph in (**B**) plots the percentage of *THR1* gene conversion-associated tetrads with a phenotypically sectored colony on SC-Thr (a non-conventional gene conversion event (post-meiotic segregation) reflecting an unrepaired mismatch at the *THR1* locus), for each of the *zip1[Δ21-163]* strains homozygous for *spo11* reduction-of-function alleles as well as the control (*SPO11+*) strain. The total number of tetrads (meioses) observed with nonmendelian segregation (conventional or non-conventional) at *THR1* is indicated (in white) at the bottom of the corresponding bar for each strain. A two-tailed *P* value for each set of mutant data compared with wild type, calculated using a Fisher’s Exact Test, is indicated above individual columns (values in red are considered statistically significant, grey not quite significant, and black not significant). All three CRISPR-Cas9 manipulated *zip1[Δ21-163]* strains display elevated frequencies of post-meiotic segregation at *THR1*, although the increase is statistically significant only in the *zip1[Δ21-163] spo11-179* strain. See Table 4 for additional detail regarding the frequency of gene conversion at *THR1* and characteristic features of post-meiotic segregation events in these strains.

**Figure S6.**
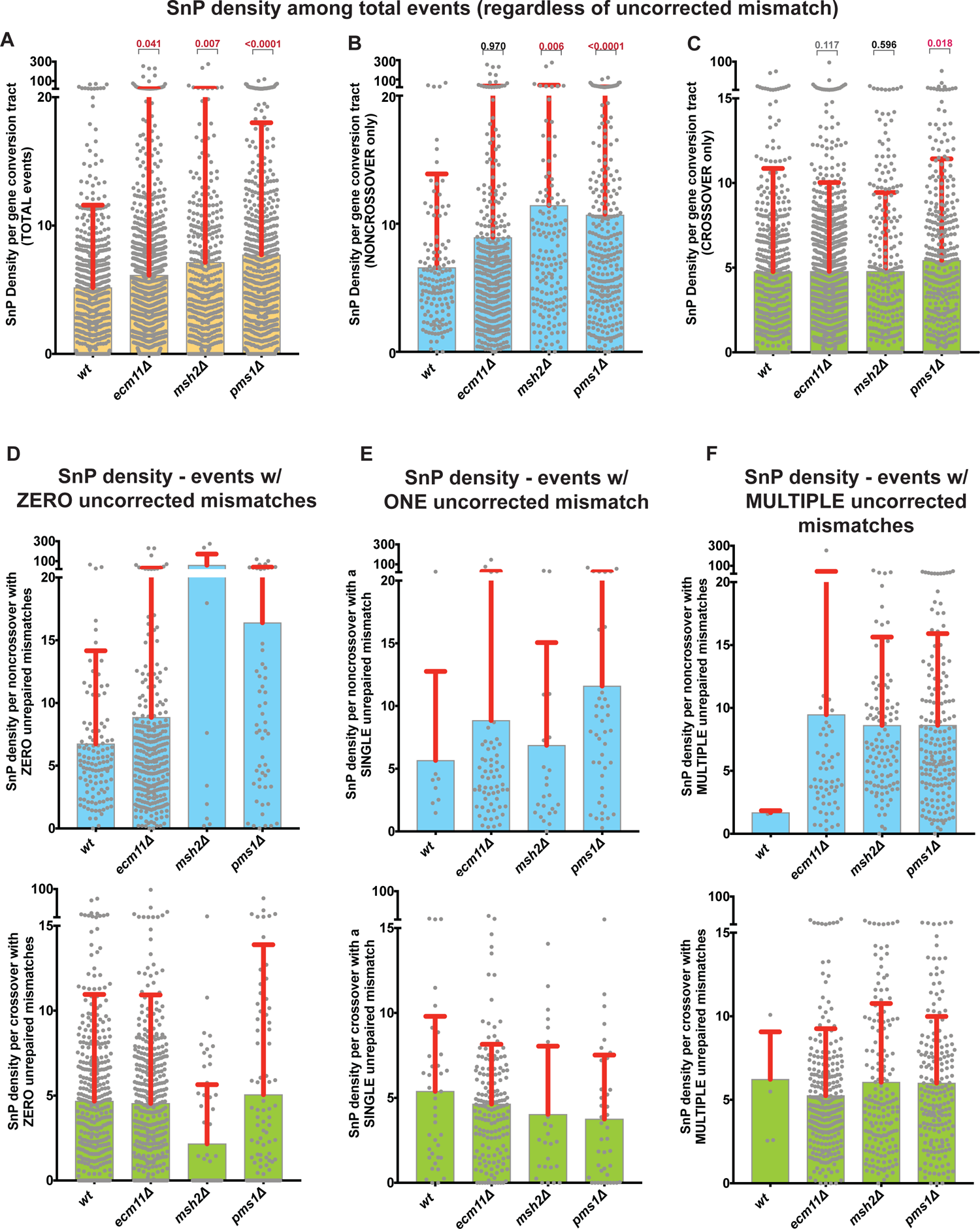
SnP density in *ecm11* gene conversion tracts is similar to wild type. Each grey circle in the scatterplots represents a single nucleotide polymorphism (SnP) density measurement for individual gene conversion tracts in four wild-type, five *ecm11*, two *msh2* and three *pms1* meioses. Graphs in (**A**-**C**) plot SnP density values for all interhomolog recombination events regardless of the presence of an uncorrected mismatch (**A**, yellow shading), for all noncrossover (E1 + E4) events regardless of an uncorrected mismatch (**B**, blue shading), and for all crossover (E2, E3, E5, E6, E7) events regardless of an uncorrected mismatch (**C**, green shading). A Mann Whitney test found little statistical difference between SnP density values for total noncrossover (**B**) or total crossover (**C**) gene conversion tracts in wild-type versus *ecm11* (although a significant difference was found when comparing total events (**A**); two-tailed *P* value indicated above mutant columns). The Mann Whitney test revealed that SnP density values associated with *msh2* and *pms1* noncrossover gene conversion tracts and *pms1* crossover gene conversion tracts are significantly different from wild-type, but SnP density values associated with *msh2* crossover tracts are not significantly different from wild type. Graphs in (**D**-**F**) plot SnP density values of noncrossover (blue shading, upper) or crossover (green shading, lower) events that display zero (**D**), one (**E**), or more than one (**F**) uncorrected mismatch. The mean for each genotype is indicated by the height of the shaded area, the standard deviation is indicated by red bars. Note chromosome 7 recombination events for two wild-type octads were excluded from these analyses, due to chromosome 7 disomy.

**Figure S7.**
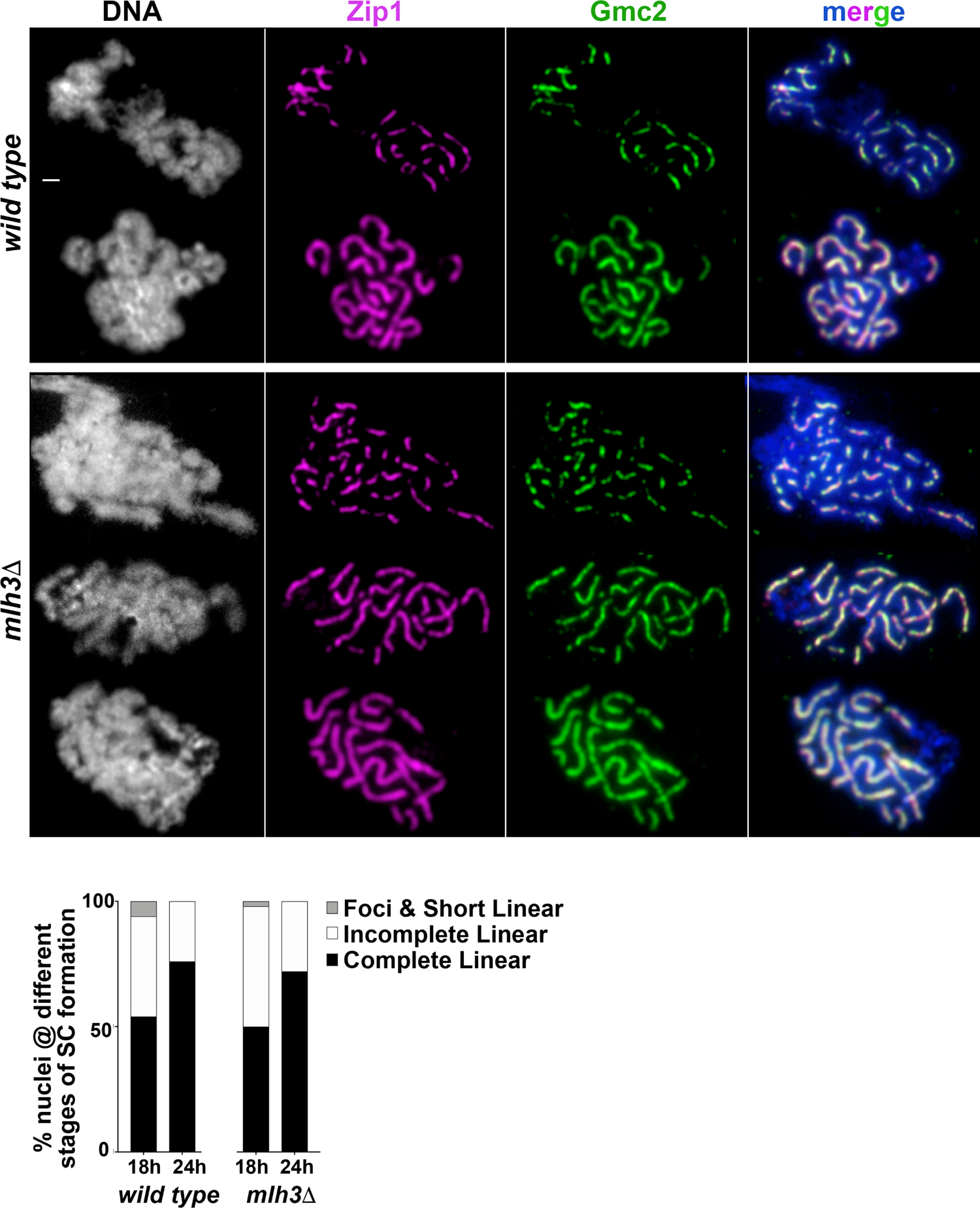
SC assembles with normal timing in *mlh3* mutant meiocytes. Images show examples of surface spread mid-meiotic prophase nuclei from *MLH3* (top two rows) and *mlh3* mutant (bottom three rows) strains. Both strains are missing the transcription factor Ndt80, to ensure that all sporulating cells, which progress through meiosis at variable rates in this strain background, halt progression at late prophase with maximal SC length. For each genotype, structures consisting of coincident Zip1 (magenta) and Gmc2 (green) were evaluated in 50 nuclei at 18 and 24 hours after placement in sporulation media. As indicated in the stacked bar graph, wild-type and *mlh3* strains at both time points displayed similar proportions of nuclei with complete SC (dark shading, second, fourth and fifth row of example images), intermediate SC (white shading, top and third row of example images) or only SC protein foci on chromatin (grey shading). Scale bar, 1 μm.

**Figure S8.**
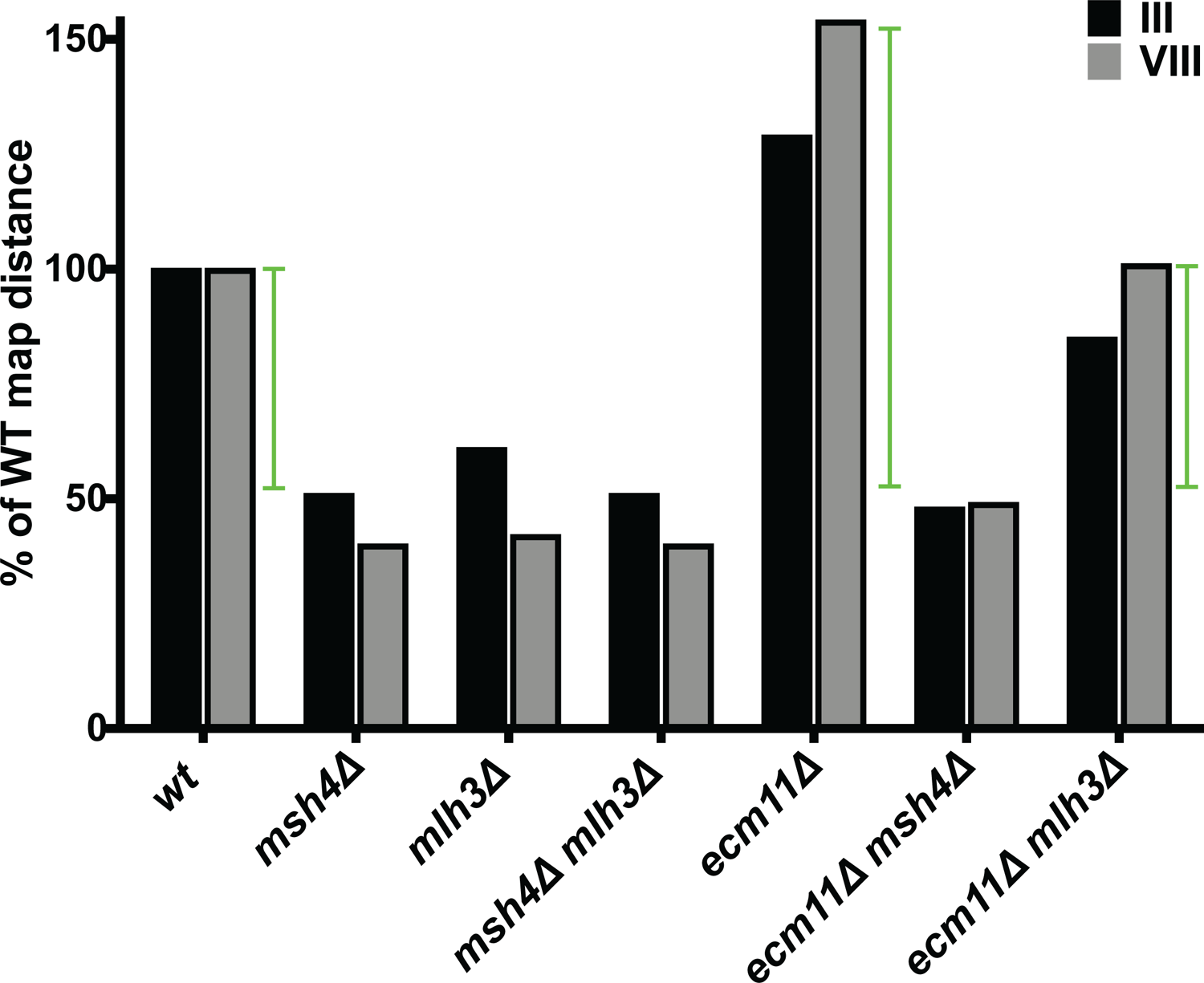
Restoration of crossovers in *mlh3* mutants by removal of Ecm11. The graph plots previously-published crossover recombination frequencies across chromosomes III (encompassing five genetic intervals) and VIII (encompassing three genetic intervals) in wild type, *mlh3, msh4 mlh3, ecm11, ecm11 msh4,* and *ecm11 mlh3* mutant strains; data were previously published in (Voelkel-Meiman *et al*. 2016) and can be found in Table S1. Crossover recombination frequencies are expressed as % of wild-type map distance, which is set at 100%.

**Table S1.**
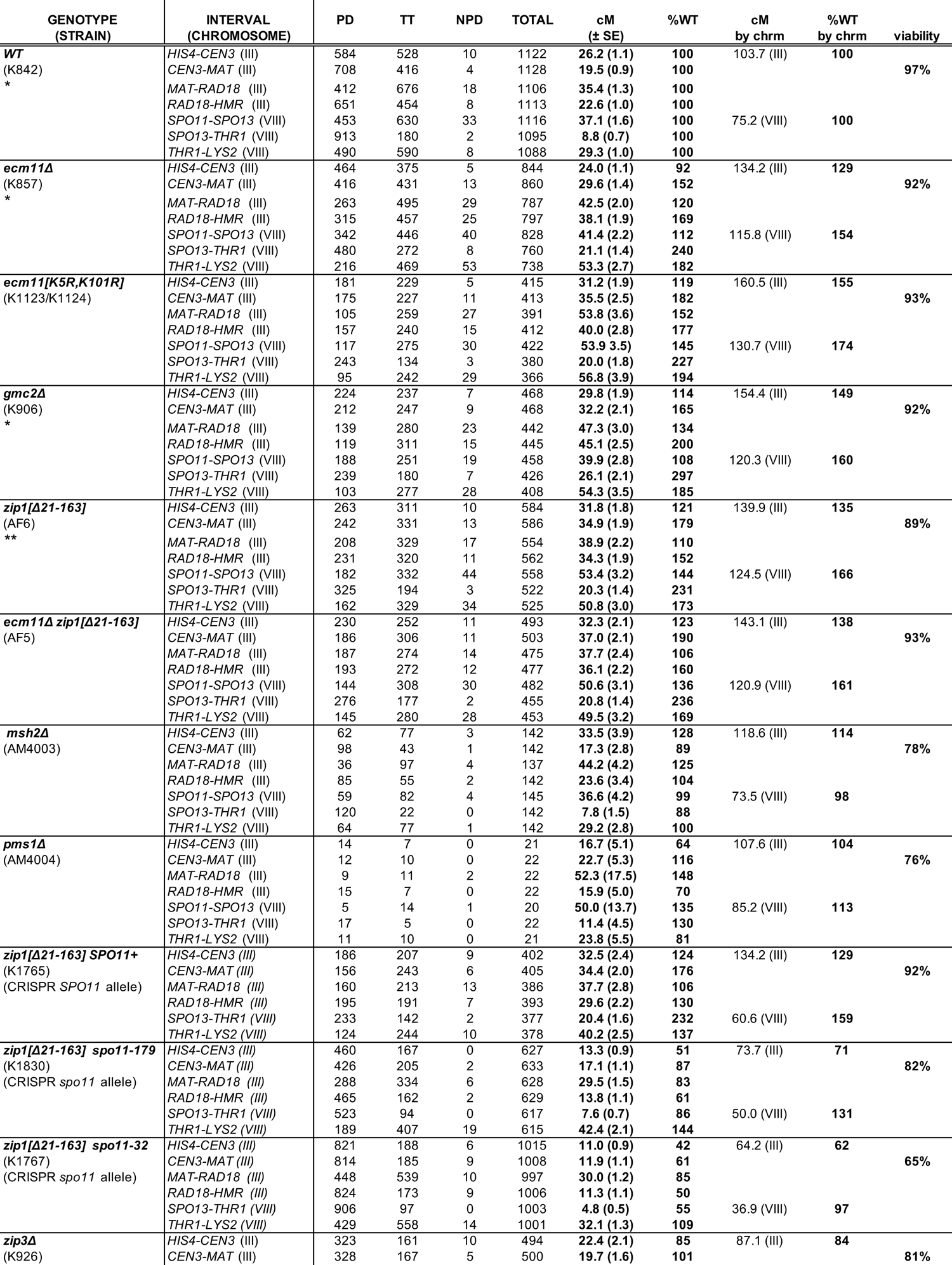
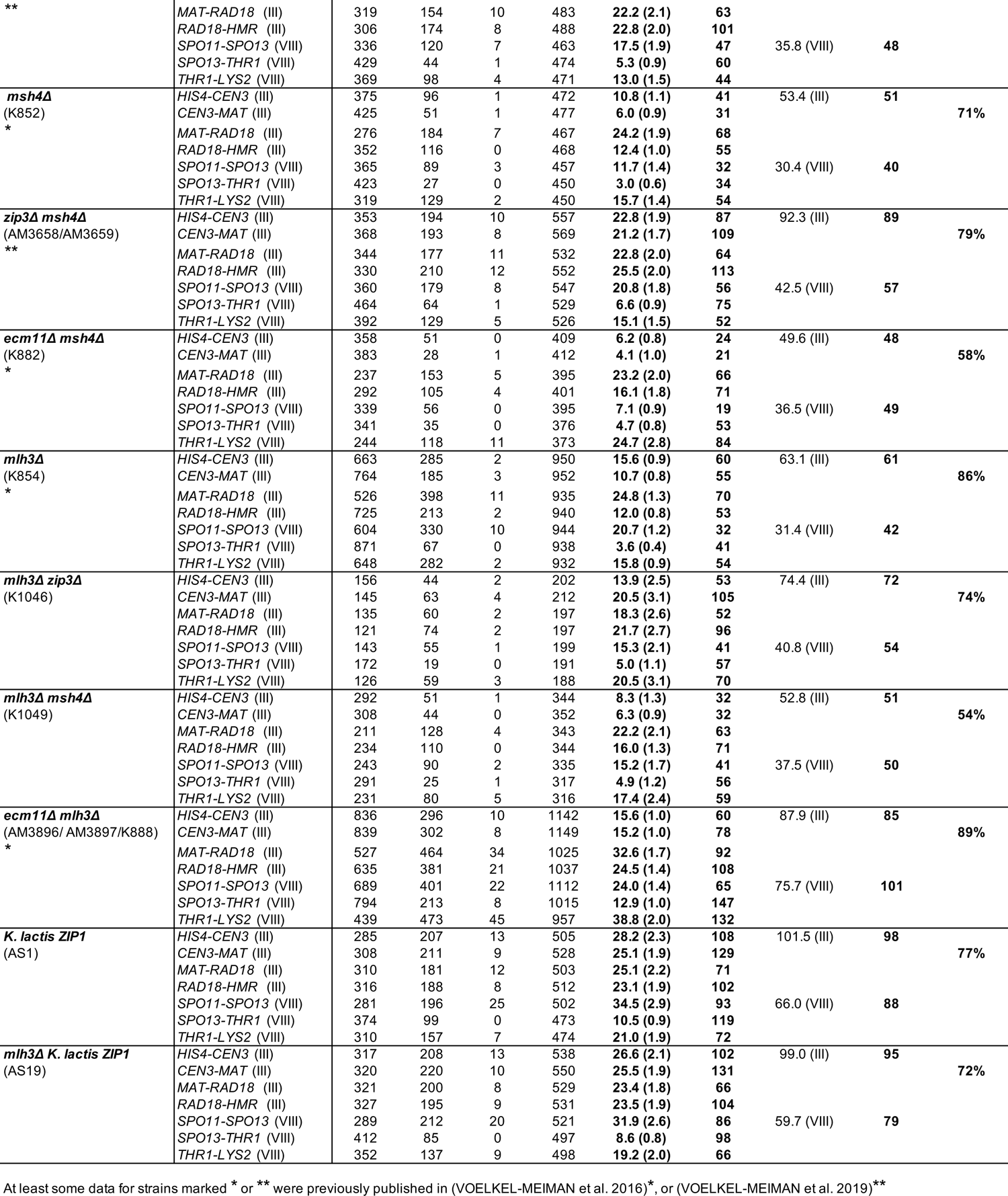
Genetic map distances of strains analyzed for post-meiotic segregation at THR1. *(Relates to Figures 1, 7, S5, S8)* Map distances were calculated using tetrad analysis as described previously (Voelkel-Meiman *et al*. 2019) in strains heterozygous for the following nine alleles: *his4-260,519, hphMX@CEN3, MAT*a*, ADE2@RAD18, natMX@HMR* (on chromosome *III*); *TRP1+@SPO11, spo13::URA3+, thr1-4, LYS2@210kb* (on chromosome VIII). Four spore-viable tetrads with no more than seven gene conversion (non-2:2) events were included in calculations. See Tables 1 and 4 for *THR1* gene conversion and post-meiotic segregation frequencies. Table indicates map distances and their corresponding percentages of the wild-type values for individual intervals, and the corresponding percentage of wild-type cumulative chromosomal map distance for chromosome III and VIII. Some data from this table is plotted in Figures S5 and S8. At least some of the data for strains marked * or ** were previously published in (Voelkel-Meiman *et al*. 2016)*, or (Voelkel-Meiman *et al*. 2019)**.

**Table S2.** Frequency of uncorrected mismatches among recombination events with different gene conversion tract lengths. *(Relates to Figure S4*) The percentage of uncorrected mismatch-carrying noncrossover (**A**) or crossover (**B**) events with specific gene conversion tract lengths. Data is consolidated from all genome-wide recombination events identified in four wild-type, five *ecm11*, two *msh2* and three *pms1* meioses (Octad Rec-Seq datasets). For each category of gene conversion tract lengths, a Fisher’s Exact Test was used to determine whether the proportion of mismatch-carrying recombination events is significantly different in any of the mutant strains relative to wild-type (*P* values are indicated on the Figure S4 graphs; red indicates statistical significance).

| **A** | % of **noncrossover** GC tracts of different lengths (n) that carry an uncorrected mismatch |  |  |  |  |  |  |
| --- | --- | --- | --- | --- | --- | --- | --- |
|  | 0-1000 bp | 1001-2000bp | 2001-3000bp | 3001-4000bp | 4001-10,000bp | 10,001-20,000bp | 20,000+bp |
| *wt* | 15.4 (26) | 4.6 (44) | 16.0 (25) | 5.0 (20) | 23.1 (13) | n/a (0) | n/a (0) |
| *ecm11* | **30.2 (96)** | **15.6 (109)** | **22.9 (70)** | **30.0 (40)** | **44.9 (98)** | **20.0 (10)** | **0 (1)** |
| *msh2* | 95.5 (66) | 97.8 (45) | 100.0 (17) | 100.0 (7) | 61.5 (13) | n/a (0) | n/a (0) |
| *pms1* | 78.4 (125) | 81.1 (74) | 87.5 (40) | 94.7 (19) | 47.1 (17) | 100.0 (1) | n/a (0) |

**Table S2. Frequency of uncorrected mismatches among recombination events with different gene conversion tract lengths.**
| **B** | % of **crossover** GC tracts of different lengths (n) that carry an uncorrected mismatch |  |  |  |  |  |  |
| --- | --- | --- | --- | --- | --- | --- | --- |
|  | 0-1000 bp | 1001-2000bp | 2001-3000bp | 3001-4000bp | 4001-10,000bp | 10,001-20,000bp | 20,000+bp |
| *wt* | 5.2 (136) | 12.0 (83) | 17.9 (56) | 13.7 (51) | 13.6 (59) | 25.0 (4) | n/a (0) |
| *ecm11* | **24.7 (186)** | **37.5 (104)** | **51.1 (88)** | **54.7 (106)** | **63.5 (342)** | **86.9 (46)** | **0 (2)** |
| *msh2* | 36.1 (97) | 91.9 (99) | 91.1 (45) | 100.0 (30) | 100.0 (38) | n/a (0) | n/a (0) |
| *pms1* | 42.2 (121) | 82.8 (87) | 78.3 (60) | 81.4 (43) | 75.8 (33) | 100.0 (1) | 0 (1) |

**Table S3.** Per-nucleotide frequency of mismatch repair errors in wild-type, ecm11, msh2, and pms1 meioses. *(Relates to Figure 5)* Table lists the per-nucleotide frequency of uncorrected mismatches identified among the sum total of noncrossover (**A**) and crossover (**B**) recombination events in four wild type, five *ecm11*, two *msh2*, and three *pms1* Octad Rec-Seq datasets. In the columns to the left of the vertical midline, an uncorrected mismatch per-nucleotide frequency *exclusive to mismatch-carrying events* was calculated by dividing the total number of uncorrected mismatches found in a given genotype by the cumulative gene conversion tract length of events carrying an uncorrected mismatch in that genotype. In columns on the right of the midline, the mismatch per nucleotide frequency for *all events* was calculated by dividing the total number of uncorrected mismatches by the cumulative gene conversion tract length of all events. Chi square with Yates Correction analysis found that mismatch repair errors are significantly more frequent among both noncrossover and crossover events in *ecm11* relative to wild-type. However, among those select recombination events susceptible to inefficient mismatch repair, the unrepaired mismatch frequency is not significantly different between *ecm11* and wild-type (according to a Fisher’s Exact Test or Chi square with Yates Correction). Note chromosome 7 recombination events for two wild-type octads were excluded from these analyses, due to chromosome 7 disomy.

| **A NONCROSSOVER EVENTS** |  |  |  |  |  |  |  |
| --- | --- | --- | --- | --- | --- | --- | --- |
|  | **Uncorrected mismatch - containing GC tract length (cumulative bp)** | **Total uncorrected mismatches among all events** | **Uncorrected mismatches per bp of GC tract, for mismatch-carrying events** | **Fishers Exact, two sided *P*** | **Mismatch-free GC tract length (cumulative bp)** | **Uncorrected mismatches per bp of GC tract, for all events** | **Chi square with Yates Correction, two sided *P*** |
| *wild type* | 35309 | 17 | 4.81E-04 |  | 243858 | 6.09E-05 |  |
| *ecm11* | 416823 | 255 | 6.12E-04 | 0.428 | 844793 | 2.02E-04 | **<0.0001** |
| *msh2* | 207732 | 802 | 3.86E-03 | **<0.0001** | 30807 | 3.36E-03 | **<0.0001** |
| *pms1* | 345315 | 1194 | 3.46E-03 | **<0.0001** | 99791 | 2.68E-03 | **<0.0001** |

**Table S3. Per-nucleotide frequency of mismatch repair errors in wild-type, *ecm11*, *msh2*, and *pms1* meioses.**
| **B CROSSOVER EVENTS** |  |  |  |  |  |  |  |
| --- | --- | --- | --- | --- | --- | --- | --- |
|  | **Uncorrected mismatch - containing GC tract length (cumulative bp)** | **Total mismatches among all events** | **Uncorrected mismatches per bp of GC tract, for mismatch-carrying events** | **Chi square with Yates Correction, two sided *P*** | **Mismatch-free GC tract length (cumulative bp)** | **Uncorrected mismatches per bp of GC tract, for all events** | **Chi square with Yates Correction, two sided *P*** |
| *wild type* | 127351 | 59 | 4.63E-04 |  | 789499 | 6.44E-05 |  |
| *ecm11* | 2270528 | 1214 | 5.35E-04 | 0.311 | 1322763 | 3.38E-04 | **<0.0001** |
| *msh2* | 494207 | 1761 | 3.56E-03 | **<0.0001** | 37193 | 3.31E-03 | **<0.0001** |
| *pms1* | 605311 | 1629 | 2.69E-03 | **<0.0001** | 145097 | 2.17E-03 | **<0.0001** |

**Table S4.**
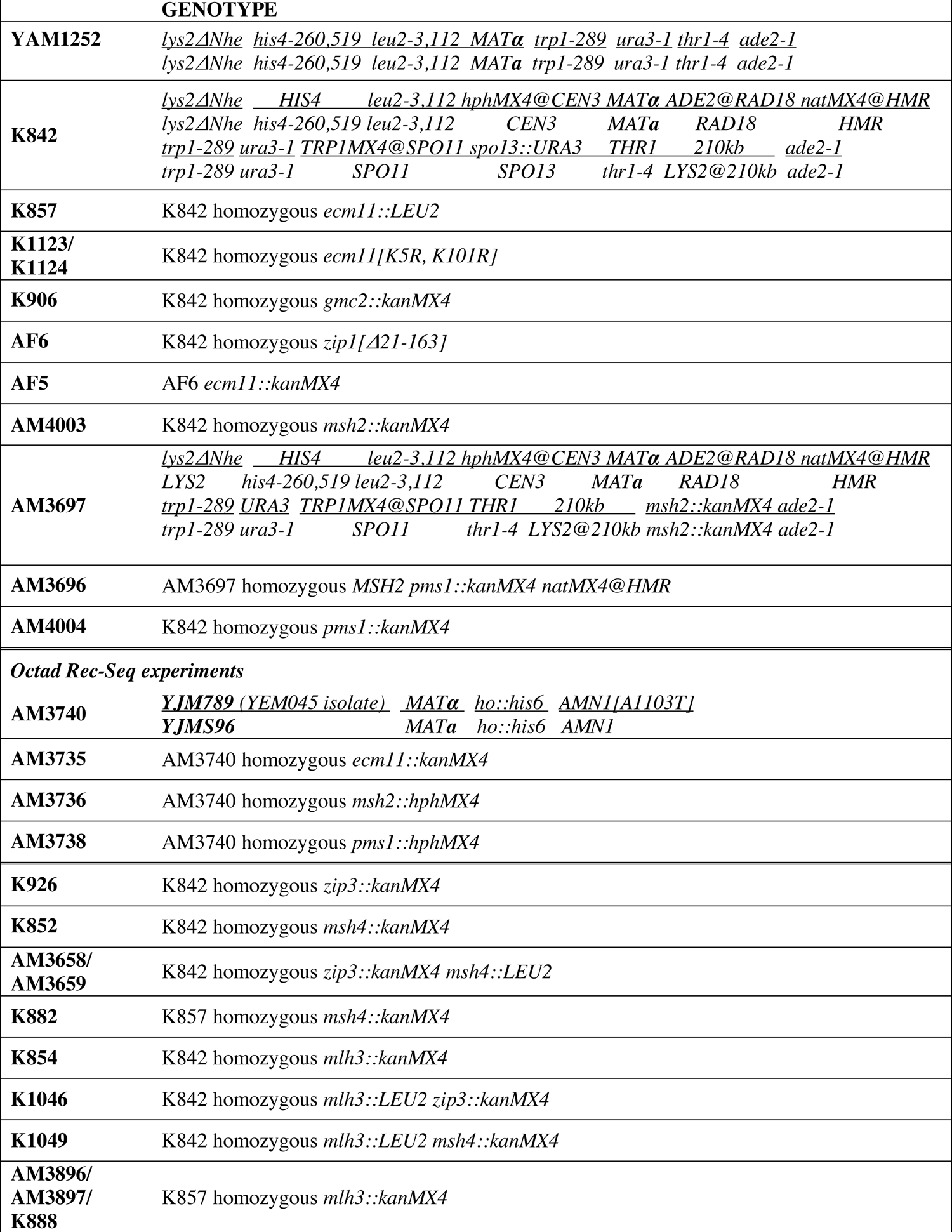
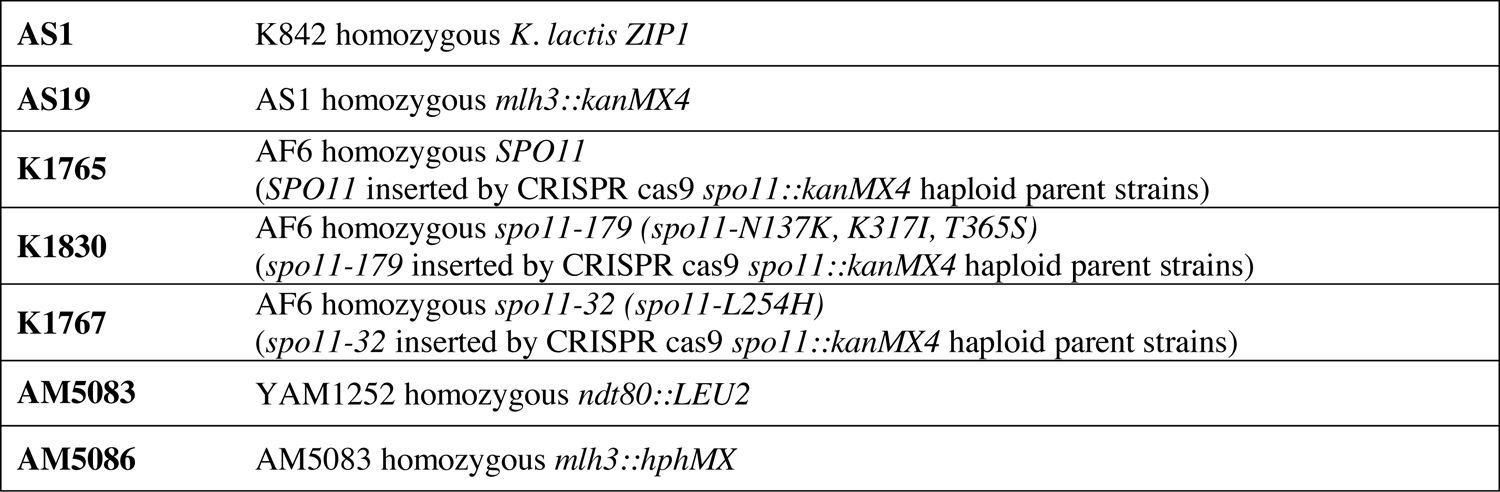
Strains used in this study. Strains are of the BR1919-8B background (Rockmill and Roeder 1998), except for those listed under the heading “Octad Rec Seq Experiments”, which are derived from a cross between YJM789 and S96 genetic backgrounds (Wei *et al*. 2007; Anderson *et al*. 2011).

## Notes

### Competing Interest Statement

The authors have declared no competing interest.

